# RNA structures within Venezuelan equine encephalitis virus E1 alter macrophage replication fitness and contribute to viral emergence

**DOI:** 10.1101/2024.04.09.588743

**Authors:** Sarah E. Hickson, Jennifer L. Hyde

## Abstract

Venezuelan equine encephalitis virus (VEEV) is a mosquito-borne +ssRNA virus belonging to the *Togaviridae*. VEEV is found throughout Central and South America and is responsible for periodic epidemic/epizootic outbreaks of febrile and encephalitic disease in equines and humans. Endemic/enzootic VEEV is transmitted between Culex mosquitoes and sylvatic rodents, whereas epidemic/epizootic VEEV is transmitted between mosquitoes and equids, which serve as amplification hosts during outbreaks. Epizootic VEEV emergence has been shown to arise from mutation of enzootic VEEV strains. Specifically, epizootic VEEV has been shown to acquire amino acid mutations in the E2 viral glycoprotein that facilitate viral entry and equine amplification. However, the abundance of synonymous mutations which accumulate across the epizootic VEEV genome suggests that other viral determinants such as RNA secondary structure may also play a role in VEEV emergence. In this study we identify novel RNA structures in the E1 gene which specifically alter replication fitness of epizootic VEEV in macrophages but not other cell types. We show that SNPs are conserved within epizootic lineages and that RNA structures are conserved across different lineages. We also identified several novel RNA-binding proteins that are necessary for altered macrophage replication. These results suggest that emergence of VEEV in nature requires multiple mutations across the viral genome, some of which alter cell-type specific replication fitness in an RNA structure-dependent manner.

**AUTHOR SUMMARY:** Understanding how viral pathogens emerge is critical for ongoing surveillance and outbreak preparedness. However, our understanding of the molecular mechanisms that drive viral emergence are still not completely understood. Emergence of the mosquito-borne virus Venezuelan equine encephalitis virus (VEEV) is known to require mutations in the viral attachment protein (E2), which drive viremia and transmission. We have observed that emergent strains (epizootic VEEV) also accumulate many silent mutations, suggesting that other determinants independent of protein sequence also contributes to emergence. In this study we identify novel RNA secondary structures associated with epizootic VEEV that alters viral replication in a cell-type dependent manner. We show that these RNA structures are conserved across epizootic viruses and identify host proteins that specifically bind these RNAs. These findings imply that viral emergence requires multiple mutations, a number of which likely alter viral structure in a manner that benefits viral replication and transmission.

## INTRODUCTION

Alphaviruses are a group of enveloped positive-sense RNA (+ssRNA) viruses belonging to the *Togaviridae* family. These viruses are transmitted by arthropod vectors and are etiological agents of several significant human and veterinary diseases. Alphaviruses are globally distributed and can be broadly classified in two groups based on their associated pathologies, chiefly arthritogenic or encephalitic. Venezuelan equine encephalitis virus (VEEV) causes periodic outbreaks of febrile and encephalitic disease in equids and humans throughout Central and South America [1]. Endemic/enzootic VEEV is predominantly transmitted between *Culex (Melanoconion)* spp. mosquitoes and sylvatic rodents such as cotton rats and spiny rats which are believed to be the major reservoir host for these endemic/enzootic viruses (subtypes ID and IE) [2]. Emergence of epidemic/epizootic VEEV (subtypes IAB and IC) occurs de novo via mutation of enzootic subtypes [3]. In contrast to endemic/enzootic VEEV, epidemic/epizootic subtypes are primarily transmitted between several mammalophilic mosquitoes and equines which are the major amplification hosts during these outbreaks [4, 5]. Spillover infections into humans also occur during epidemic/epizootic episodes and can be associated with severe encephalitic disease and death, as well as long-term debilitating sequelae [6]. Repeated emergence of epidemic/epizootic VEEV has previously been shown to involve mutation of the viral attachment protein (E2) of ID endemic/enzootic subtypes which give rise to epidemic/epizootic VEEV subtypes IAB and IC [7–9]. E2 mutations were found to facilitate increased replication levels in horses, heightened virulence, and adaptation to epizootic mosquito vectors [4, 10]. While these mutations alone have been demonstrated to be sufficient for imparting epizootic phenotypes in a laboratory setting, epidemic/epizootic subtypes contain numerous additional mutations across the viral genome which studies suggest may contribute to epidemic/epizootic emergence [11, 12].

The VEEV genome is approximately 11.5kb in length and contains a 5’methylguanosine (m7G) cap and a 3’ polyA tail [13]. The genome consists of two open reading frames, ORF1 which encodes four non-structural proteins (nsp1-4) and ORF2 which encodes a subgenomic RNA from which the viral structural proteins are translated. We have previously shown that RNA structures present in the 5’UTR of the VEEV and Sindbis virus (SINV) confer resistance to the interferon stimulated gene (ISG) IFIT1, by preventing IFIT1 recognition of viral m7G capped RNA [14]. Similarly, we have observed that changes in VEEV 3’UTR structure alters IFIT2-mediated restriction of viral replication in a subtype-dependent manner [15]. Notably, most SNPs acquired by epidemic/epizootic strains following VEEV emergence are synonymous, suggesting that in addition to protein coding mutations in E2, changes in viral RNA structure may contribute to emergence of epidemic/epizootic VEEV. In this study we identify novel RNA structures in E1 that alter replication in macrophages which are early targets of VEEV infection in vivo. Conservation of SNPs and RNA secondary structures in this region suggest that these structures may contribute to emergence of epidemic/epizootic VEEV. These findings have significance for our understanding of VEEV evolution and emergence.

## RESULTS

To first identify putative RNA structures that differ between endemic/enzootic and epidemic/epizootic strains we used phylogenetic analysis to identify closely related pairs of enzootic and epizootic strains for subsequent RNA structure analysis (**Extended Data Fig. 1**). We compared 143 isolates and identified three enzootic (subtype ID) strains (R16905, 307537, and 204381) which exhibited 99.4%, 96.5%, and 96.2% sequence identity to the epizootic (subtype IAB) vaccine strain TC83. For downstream RNA structure analysis, we chose to compare TC83 and 307537. TC83 is a BSL2 attenuated vaccine strain developed by serial passage of strain Trinidad donkey (TRD) which was originally isolated from a sick donkey during an epizootic outbreak in Trinidad [13, 16]. TC83 shows 99.9% sequence identity with TRD but contains attenuating mutations in the 5’UTR and the viral attachment protein E2 [13, 16]. 307537 is a geographically distinct strain first isolated from mosquitoes and shares 96.5% sequence identity with TC83. To determine the predicted secondary structure of each viral genome, we used RNAfold [17, 18] [19] [20] to perform a sliding window analysis of each strain and generate an RNA structure score (RSS) for each window (**Fig. 1B**). The RSS is generated by dividing the frequency of the minimum free energy structure (MFE) by the ensemble diversity (ED), and thus captures some qualitative data of RNA secondary structures formed by that sequence. In this instance, a higher RSS suggests the presence of RNA structures which are more thermodynamically stable and have a higher probability of forming. By reducing the complexity of RNA secondary structure to a single numerical value, we can compare large groups of sequences (e.g. phylogenetic analysis) and identify RNA ‘signatures’ which may be unique or conserved within these groups. Our analysis revealed several regions with highly stable putative RNA structures (z-score >2), including nsp1, nsp2, nsp4, capsid, and E1 (**Fig. 1B, Extended Data Fig. 2**). Previously defined functionally relevant RNA structures were also identified using this analysis, notably the nsp1 packaging signal [21], and the ribosomal frameshift (RFS) motif in 6K/E1 which is required for production of TF protein [22, 23]. In addition, we identified several regions in which the predicted RNA structure differed between TC83 and 307537, including within E1 (**Fig. 1C**). As we observed a high proportion of synonymous mutations in this gene (97.6%; **Fig. 1A**) and have previously shown that RNA structures proximal to this gene (3’UTR) alter replication properties of VEEV [15], we sought to define the role of E1 RNA structures in viral replication and their potential contribution to emergence of epizootic VEEV.

**Figure 1.**
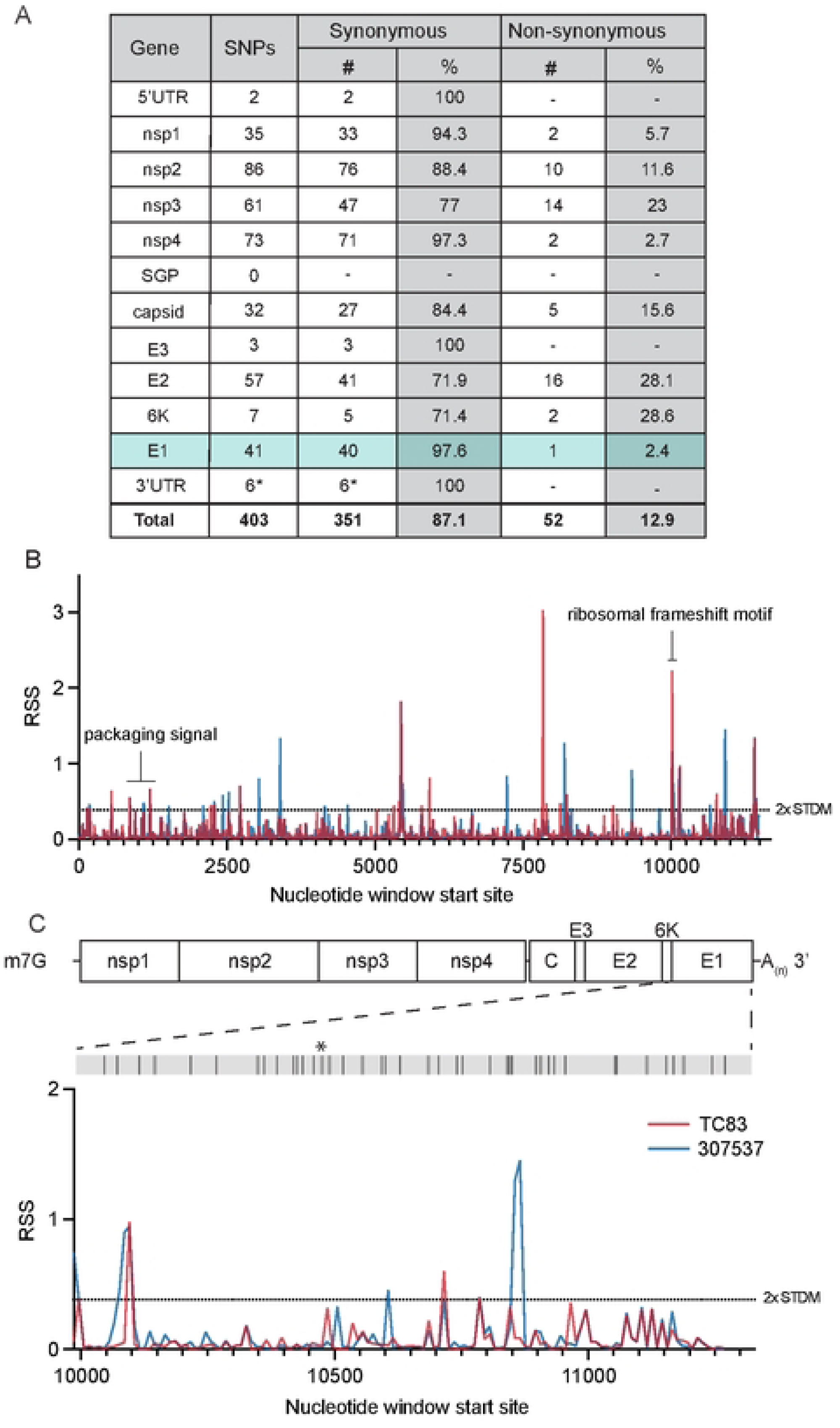
Predicted RNA secondary structure of E1 differs between epizootic and enzootic VEEV. **(A)** Summary of all SNPs identified between TC83 and 307537. **(B)** RNA structure analysis of subtype IAB and ID VEEV. RNA structure prediction of VEEV strain TC83 (IAB; accession L01443) and strain 307537 (ID; accession KC344519) was performed using RNAfold [17] (window size = 50nt, step size = 10 nt). The RNA structure score (RSS; frequency of MFE/ensemble diversity) is plotted against the nt window start site. Higher RSS indicates greater thermodynamic stability of predicted structures. The 2-fold standard deviation is indicated by a dotted line. **(C)** RSS analysis of gene E1 from strains TC83 and 307537. Location of all SNPs across E1, including a single coding change (*), are depicted in the grey bar above.

To determine whether changes in predicted E1 RNA structures alter VEEV replication properties, we generated a chimeric TC83 virus encoding all synonymous changes from E1 of strain 307537 (TC83/E1_IDsyn_) (**Fig. 2A**). To disentangle confounding effects of amino acid changes on replication phenotypes, this chimera excluded the single protein coding mutation found within this region (nucleotide (nt) 10,481, **Fig. 1C**). Notably, inclusion of this mutation in our structure analysis did not significantly alter the RSS in this region, and thus was predicted to have minimal effect on E1 RNA structure (**Fig. 1C**). We then compared replication kinetics of TC83 and TC83/E1_IDsyn_ in several cell types including the macrophage cell line Raw264.7, primary bone marrow-derived macrophages (BMDM), primary bone marrow derived dendritic cells (BMDC), and mouse embryonic fibroblasts (MEF) **(Fig. 2B-E**). Here, cells were infected with WT or mutant viruses at an MOI of 0.1 and production of infectious virus measured over time by focus forming assay (FFA). Myeloid cells including macrophages are early targets of encephalitic alphavirus infection in vivo and have been shown to be a source of type I IFN production early during infection [24, 25]. Thus, replication fitness in macrophages would be predicted to have significant impacts on outcomes of VEEV infection *in vivo*. In both Raw264.7 and primary BMDM we observed an increase in TC83/E1_IDsyn_ relative to TC83 (at 12hpi, 8-fold in Raw264.7, *P* = 0.0035; 10-fold in BMDM, *P* = 0.0005) (**Fig. 2B, C**). Remarkably, we observed no significant difference in replication of TC83 and TC83/E1_IDsyn_ in either BMDC or MEF (**Fig. 2D, E**), indicating that RNA sequences from E1 of enzootic VEEV specifically increases replication fitness in macrophages but not in other cell types.

**Figure 2.**
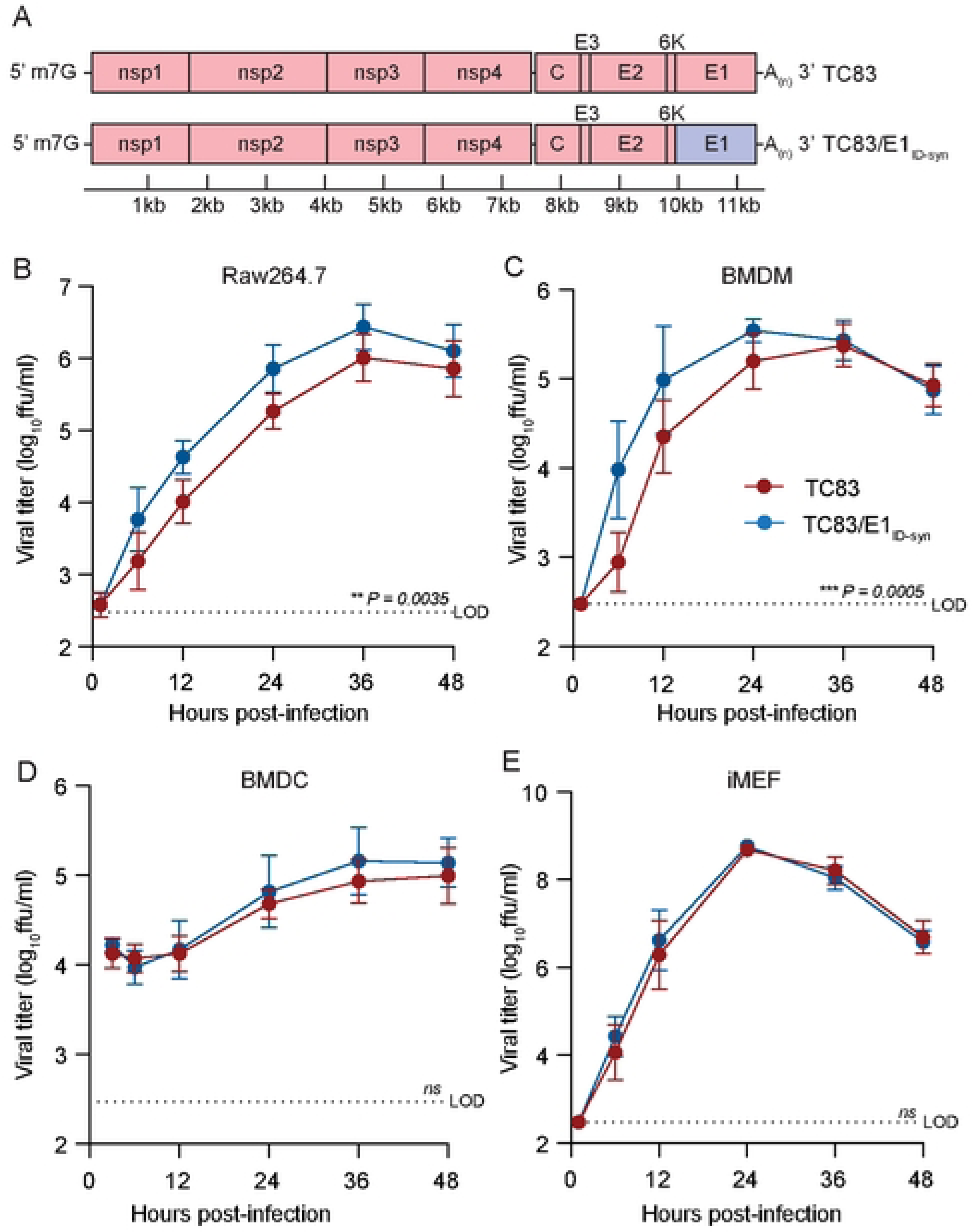
Changes in E1 RNA sequence alters viral replication fitness in macrophages, but not other cell types. **(A)**. Schematic representation of the TC83 and TC83 E1 RNA mutant (TC83/E1_ID-syn_) genomes. Synonymous SNPs from E1 of enzootic VEEV (strain 307537) were introduced into the vaccine epizootic VEEV (strain TC83) to generate an RNA mutant (TC83/E1_ID-syn_). The single coding change present in E1 (Fig 1C, asterisk) was omitted from the mutant. Red denotes sequences from the parent epizootic strain (TC83), and blue denotes sequences from the enzootic strain (307537). **(B-E)**. Replication kinetics of VEEV TC83 and TC83/E1_ID-syn_ in **(B)** Raw264.7, **(C)** primary bone marrow-derived macrophages (BMDMs), **(D)** primary bone marrow-derived dendritic cells (BMDCs), and **(E)** immortalized mouse embryonic fibroblasts (iMEF). Cells were infected with indicated viruses at a MOI of 0.1 (Raw264.5, BMDMs, iMEF) or MOI 0.01 (BMDCs). Cell culture supernatant was serially harvested at 1, 6, 12, 24, 36, and 48 hpi and infectious virus was titered using focus forming assay (FFA). Each experiment was performed in triplicate three to four times independently and the mean and SD are graphed. Statistical analysis was performed by calculating the area under the curve (AUC) for each replicate, and the AUC values from WT and mutant viruses were analyzed by unpaired t-test.

Type-I IFN is important in restricting replication and pathogenesis of alphaviruses [26–28], and we have previously shown that VEEV RNA structure facilitates evasion of IFN-stimulated genes (ISGs) [14, 15]. Thus, we hypothesized that putative E1 RNA structures from TC83/E1_IDsyn_ could enhance replication in macrophages by facilitating evasion of host antiviral immunity. Specifically, we predicted that mutant E1 RNAs may evade sensing of VEEV RNA by RLRs RIG-I and MDA-5 which are known to play a role in alphavirus RNA sensing, particularly of 3’ RNAs [29, 30]. To test this hypothesis, we used CRISPR to generate *Ddx58* and *Ifih1* knock out (KO) Raw264.7 macrophages and compared replication kinetics of TC83 and TC83/E1_ID-syn_ in these cells (**Fig. 3A-C; Extended Data Fig. 3**). Contrary to our expectations, TC83 replication was still impaired relative to TC83/E1_ID-syn_ in both the absence and presence of RIG-I or MDA-5 expression (**Fig. 3A-C**). To confirm these data, we used transient siRNA knock down of *Ddx58* and *Ifih1*, as well as *Irf3* (**Fig. 3D; Extended Data Fig. 3**) and examined titers of TC83 and TC83/E1_ID-syn_ compared to cells treated with a non-silencing control (NSC) siRNA. We predicted that if enhanced replication of TC83/E1_IDsyn_ was due to evasion of RLR-dependent sensing and antiviral restriction then knock down of RLR expression or expression of downstream signaling molecules (IRF3) would result in an increase in replication of TC83 but no change in the replication of TC83/E1_ID-syn_. However, consistent with CRISPR data, we observed no increase in replication of TC83 in the absence of either RLR expression or IRF3. Furthermore, knockdown of *Irf3* did not lead to an increase in TC83 replication relative to TC83/E1_ID-syn_, suggesting that preferential sensing and/or inhibition of TC83 RNA cannot explain the observed replication differences in macrophages.

**Figure 3.**
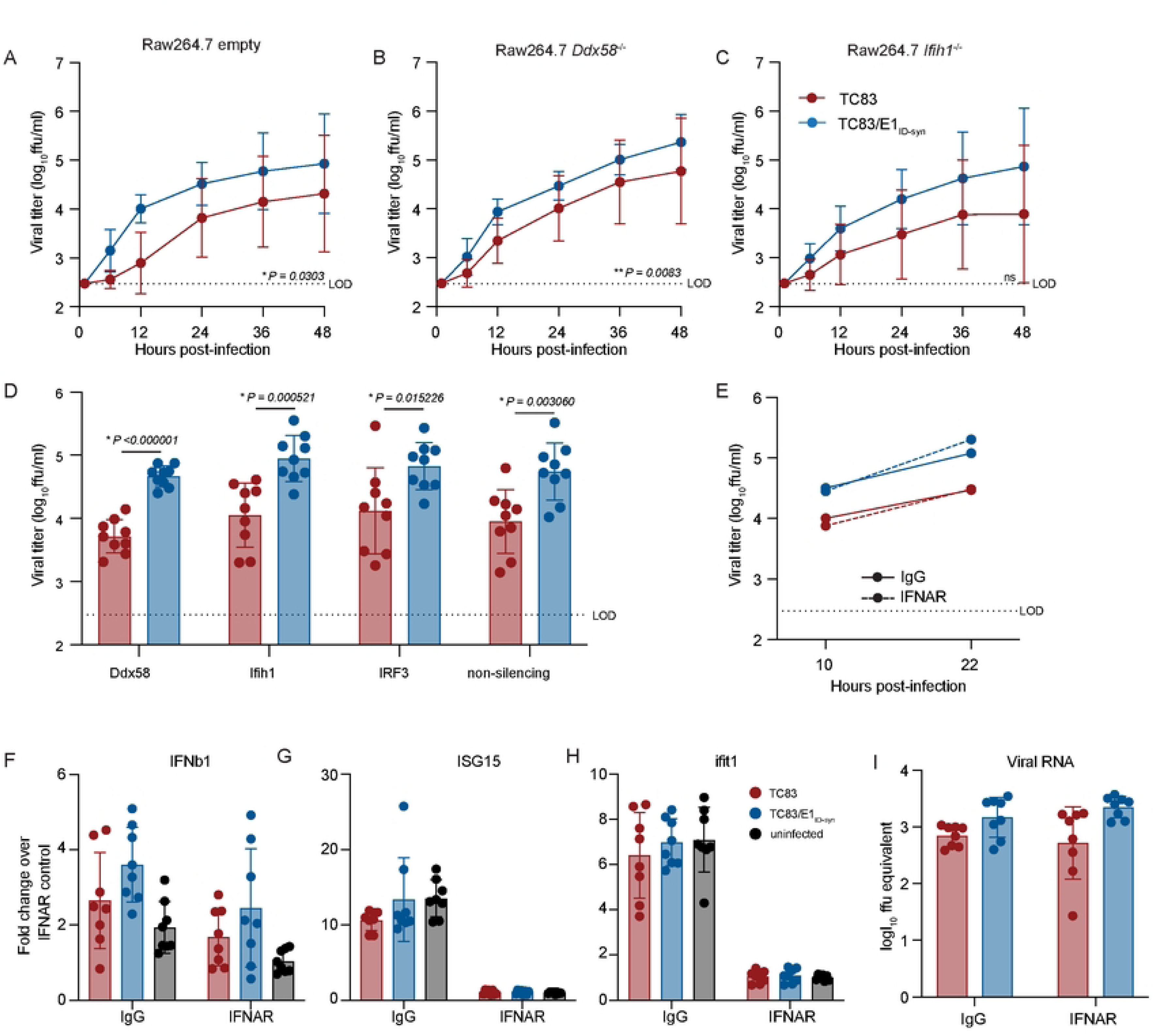
Differential macrophage replication of TC83 and TC83/E1_ID-syn_ viruses is IFN- and RLR-independent. **(A-C)** Replication kinetics of VEEV TC83 and TC83/E1_ID-syn_ in **(A)** empty vector, **(B)** *Ddx58*^-/-^, and **(C)** *Ifih1*^-/-^ CRISPR Raw264.7 cells. Cells were infected with indicated viruses at a MOI of 0.1. Cell culture supernatant was serially harvested at 1, 6, 12, 24, 36, and 48 hpi and infectious virus was titered using focus forming assay (FFA). Each experiment was performed in triplicate three times independently and the mean and SD are graphed. Statistical analysis was performed by calculating the area under the curve (AUC) for each replicate and experiment, and the AUC values for each virus analyzed by unpaired t-test. **(D)** Raw264.7 were treated with non-silecing control (NSC) siRNA or siRNA targeting *Mavs*, *Ddx58*, *Ifih1*, or *Irf3*. Cell culture supernatants were harvested at 24 hpi and infectious virus quantified by FFA. **(E)** Raw264.7 were pretreated for 1 hour with 10µg of IgG or IFNAR blocking antibody, then infected with TC83 or TC83/E1_ID-syn_ at an MOI of 0.1 in the presence of antibody. Infectious virus from cell culture supernatants harvested at 10 and 22 hpi was titered by FFA. Each experiment was performed three times independently. **(F, G)** IFNAR blocking antibody assays were performed in WT Raw264.7 as described in E, and cell lysates collected at 22 hpi. IFNb1, ISG15, Ifit1 and VEEV viral RNA transcripts quantified by qRT-PCR. Gene expression within samples was normalized to hprt, and fold change in gene expression relative to IFNAR samples was calculated. Each experiment was performed three times independently in duplicate or triplicate, and statistical analysis was performed using unpaired t-test.

While these data did not support a role of RLR-mediated RNA sensing in differential replication of TC83 and TC83/E1_ID-syn_, we could not rule out a role for other RNA sensing pathways or antiviral effectors which are independent of these pathways. As antiviral signaling pathways converge on expression of type-I IFNs which are critical for restriction of alphaviruses through expression of antiviral effectors, we examined whether differences in type-I IFN signaling and ISG expression accounted for enhanced replication of TC83/E1_ID-syn_. To determine whether infection with TC83 or TC83/E1_ID-syn_ led to differential activation of type-I IFN responses, Raw264.7 were treated with antibodies specific for the IFN-alpha receptor (IFNAR) or an IgG isotype control antibody prior to and during infection (**Fig. 3E**). We expected that if diminished TC83 replication was due to impaired evasion of RNA sensing and IFN activation then IFNAR blockade would result in an increase in viral replication to levels similar to the mutant. However, while IFNAR blockade led to a significant inhibition of ISG expression as measured by qRT-PCR (**Fig. 3F-H**), neither infectious viral titers nor viral RNA production were affected when compared to treatment with an isotype control (**Fig. 3E and I**). Collectively, this data suggests that differential replication of TC83 and TC83/E1_ID-syn_ cannot be explained by altered evasion or induction of IFN or ISG expression by either virus.

To unveil what IFN-independent mechanisms might underlie the observed differences in TC83 and TC83/E1_ID-syn_ replication (**Fig. 2**), we used a proteomics approach to identify host proteins which interact differently with TC83 and TC83/E1_ID-syn_ RNA. We hypothesized that changes in primary sequence and/or secondary structure could alter the viral RNA-protein interactome leading to changes in replication. Specifically, we predicted that antiviral RNA-binding proteins (RBPs) would be enriched for TC83 RNA or that proviral RBPs would be enriched for TC83/E1_ID-syn_ RNA. To define the RNA-protein interactome of TC83 or TC83/E1_ID-syn_, Raw264.7 were infected at MOI 0.1 and viral RNA immunoprecipitated at 24 hpi using the J2 anti-dsRNA antibody [31]. RNA-bound protein targets were then purified and identified using Liquid chromatography–mass spectrometry (LC-MS). A total of 166 proteins were identified (**Supplementary Figure/File X**). Differential enrichment of protein targets for each virus was calculated and targets prioritized as follows: (i) targets with spectral counts >10; (ii) targets showing >2-fold enrichment over the paired IgG control in at least one sample; (iii) targets showing >2-fold enrichment in TC83 vs TC83/E1_ID-syn_ (or vice versa); (iv) targets with known RBP activity (based on RBPbase and GO terms). While MS data was generated from two independent experiments (**Fig. 4**), we observed much lower spectral counts for targets in the second experiment as well as lower enrichment scores overall. Nonetheless, we identified several targets in both screens that were either differentially enriched for the WT or mutant virus (>1.5-fold; FBL, NOP58, CHTOP) or which were enriched equally for both (DHX9, ADAR, YBX1). Based on the more robust nature of the data set, downstream targets chosen for validation were based on data from experiment 1. Based on the criteria above we identified a total of 24 RBPs which showed differential binding to either the TC83 or TC83/E1_ID-syn_ genomes (**Fig. 4A**). In addition, we also identified several highly abundant targets (YBX1, HNRNPC, HNRNPM, and ADAR1) that were equally enriched for both viruses which have also been identified in previous studies as interacting with alphavirus RNA and would not be expected to be differentially enriched [32–34](**Fig. 4B**). Remarkably, with the exception of UBTF and DHX38, all identified targets were found to be enriched for the mutant, suggesting that enhanced replication of TC83/E1_ID-syn_ is not due to evasion of antiviral factors that restrict TC83, but recruitment of proviral RBPs to TC83/E1_ID-syn_. Pathway analysis of these top hits showed enrichment for RBPs associated with snoRNAs and more broadly RNA metabolism (**Fig. 4C**). To validate IP-MS findings and determine which RBPs were necessary for enhanced viral replication of TC83/E1_ID-syn_ in macrophages, we used siRNA to inhibit expression of 11 of these targets in Raw264.7 and assess replication of WT and mutant viruses in these cells (**Fig. 4D, Extended Data Fig. 4**). Here, Raw264.7 were transfected with NSC siRNA or a pool of 3 gene-specific siRNAs, infected with TC83 and TC83/E1_ID-syn_ at MOI 0.1, and infectious virus quantified from supernatants by FFA. We observed that knock down of four of these targets (*Thrap3*, *Fbl*, *Ubap2l*, and *Dhx38*) led to reduced TC83/E1_ID-syn_ replication to levels comparable to TC83, as compared to NSC-treated cells. To exclude the possibility that increased cell death following gene knock down could account for non-specific changes in viral replication in siRNA versus NSC treated cells we also measured cell viability in siRNA treated cells following infection at 24hpi (**Extended Data Fig. 4C**). Here, we observed no change, or only modest changes in cell viability which could not account for the decrease in TC83/E1_ID-syn_ replication observed.

**Figure 4.**
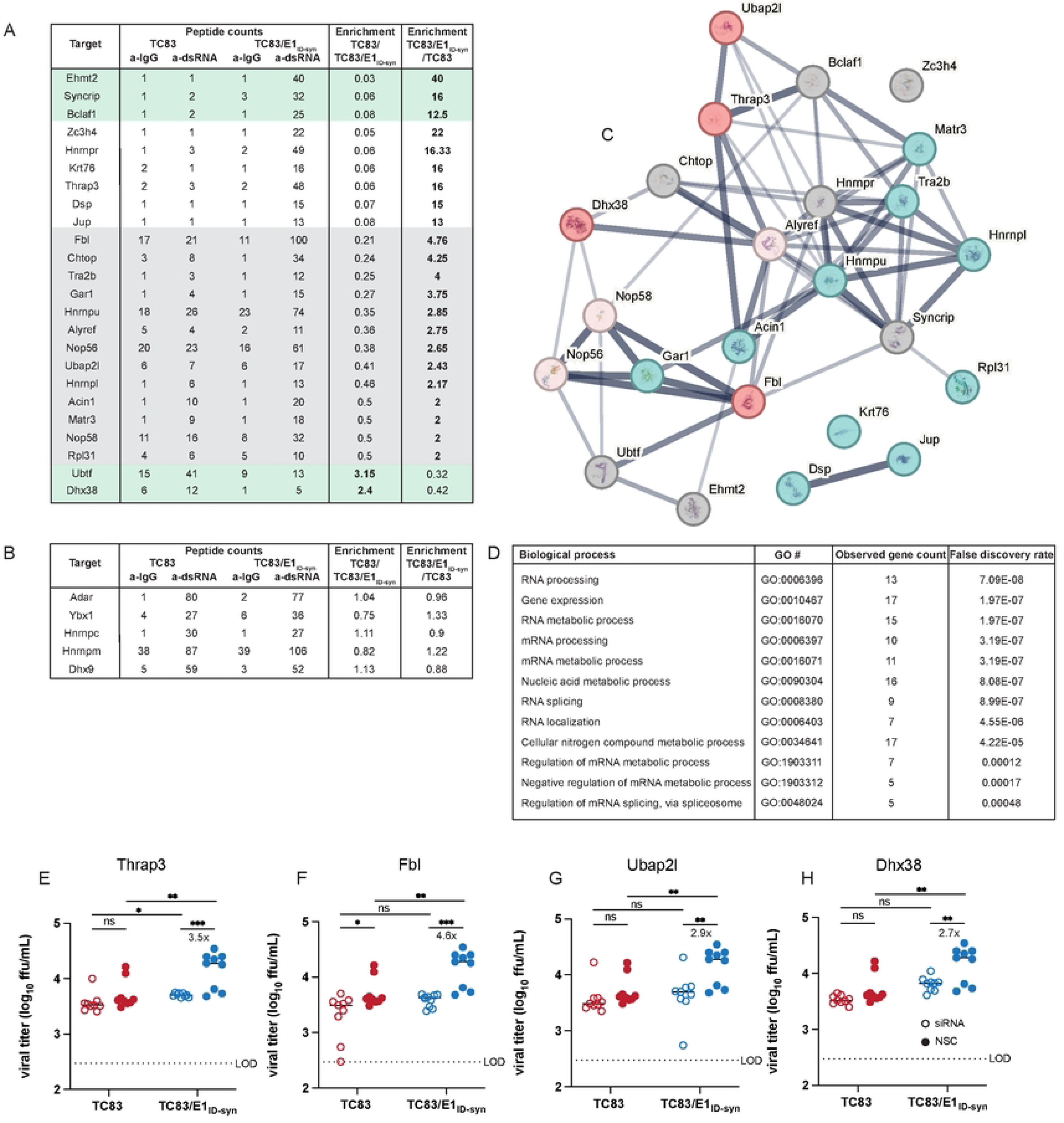
Increased macrophage replication fitness of TC83/E1_IDsyn_ is dependent on expression of RNA binding proteins Fbl, Thrap3, Ubap2l, and Dhx38. **(A)** Top hits from dsRNA immunoprecipitation-mass spectrometry (IP-MS) of TC83 and TC83/E1_IDsyn_ in Raw264.7. Raw264.7 cells were infected with TC83 or TC83/E1_IDsyn_ at an MOI of 0.1, and viral dsRNA isolated from lysates at 24 hpi using J2 dsRNA antibody [31] or IgG isotype control. RNA-bound proteins were identified by MS, and fold-enrichment of spectral counts relative to IgG controls was calculated. Prioritized hits were chosen based on fold enrichment scores, total spectral counts, and whether targets are known RNA binding proteins (RBPbase hits). **(B)** Hits equally enriched in TC83 and TC83/E1_IDsyn_. **(C)** STRING network analysis of top proteomics hits. Candidates meeting the cutoff criteria (A) were subjected to Protein-Protein Interaction Networks Functional Enrichment Analysis. Candidate proteins identified in the screen are highlighted in red and interacting proteins in blue. **(D) Enriched biological process GO terms that with a p-value >0.001, along with the observed gene count present in the STRING network (E-H)** Raw264.7 were transfected with control (NSC) or pooled (3 siRNA) gene specific siRNAs targeting **(E)** Thrap3, **(F)** Fbl, **(G)** Ubap2l, or **(H)** Dhx38 for 24 hours. Cells were infected TC83 or TC83/E1_ID-syn_ at an MOI of 0.1, cell culture supernatants collected at 24hpi, and infectious virus titered by FFA. All siRNAs were assayed simultaneously but for visual clarity, data for each gene is shown separately along with the shared control siRNA samples. Each experiment was performed in triplicate three times independently and the mean and SD are graphed. Statistical analysis was performed using unpaired t-test. ** >0.001, ***>0.0001. Fold change and p-values are indicated on each graph.

Analysis of primary E1 sequences from TC83 and TC83/E1_ID-syn_ failed to reveal obvious recognition motifs for any of the targets identified in our proteomics study. Thus, we generated additional E1 mutants to map regions within E1 necessary for differential macrophage replication and RBP recruitment (**Fig. 5A**). Here, Raw264.7 cells were infected with parent or mutant viruses at MOI of 0.1 and infectious titers at 12 and 24hpi assessed by FFA (**Fig. 5B, C**). We initially compared replication of two mutants in which the 5’ or 3’ half of E1 was exchanged between TC83 and TC83/E1_ID-syn_ (mutant 1 and 2). Surprisingly, both mutant 1 and 2 replicated identically to the parent TC83 virus, suggesting that the element responsible for differential replication was located in the middle of E1 and was disrupted in these two mutants. To test this, we generated another mutant (mutant 3) which contained only SNPs from the central region of TC83/E1_ID-syn_ E1 (nts 10,466-10,843) and compared replication of all viruses in Raw264.7 (**Fig. 5B, C**). In contrast to mutant 1 and 2, mutant 3 replicated to similar levels as that of TC83/E1_ID-syn_, confirming that the elements responsible for enhanced macrophage replication are located in the central region of E1.

**Figure 5.**
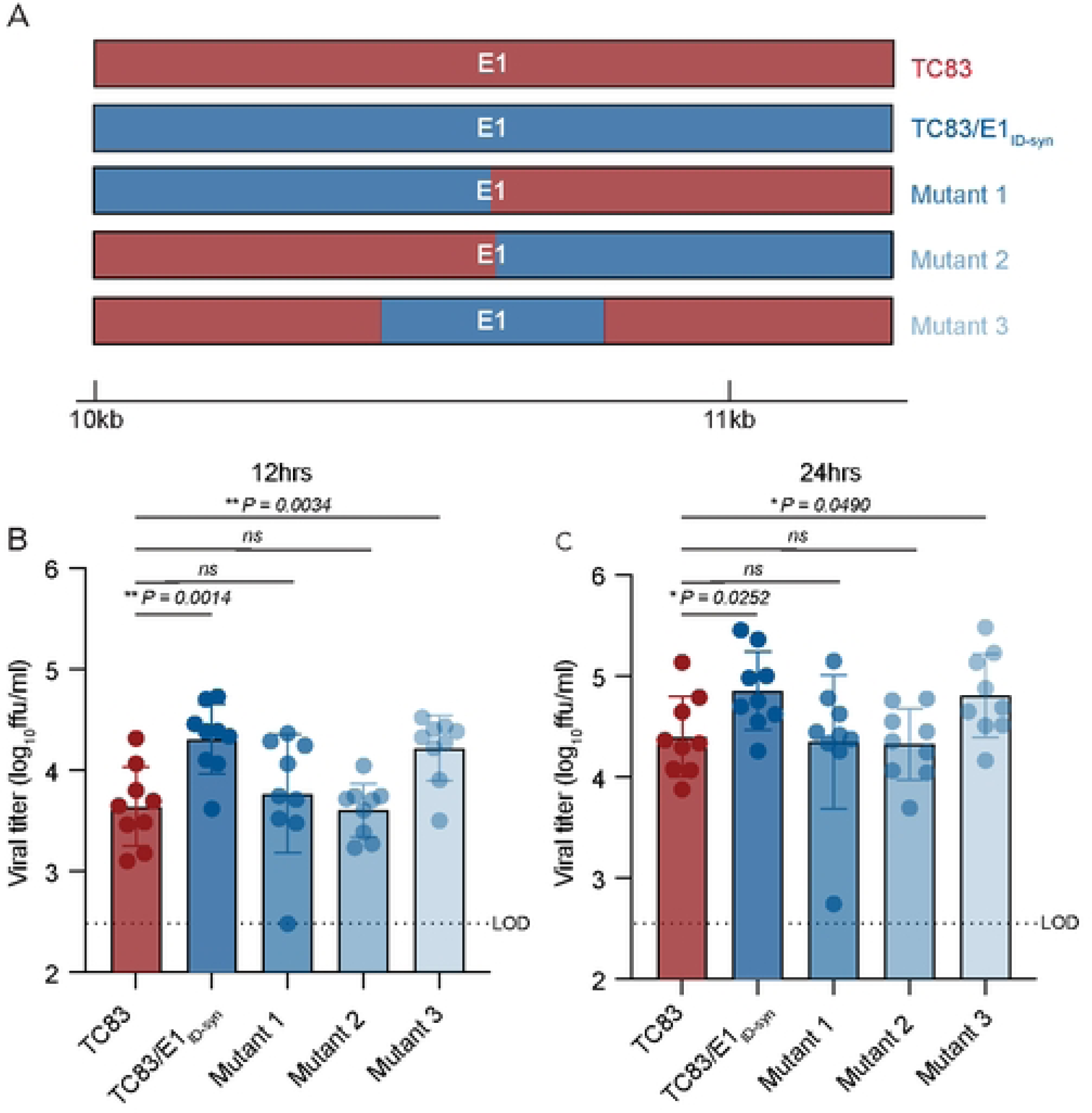
RNA sequences in the central domain of E1 enhance macrophage replication of TC83/E1_IDsyn_. **(A)** Schematic representation of mutant viruses constructed. RNA chimeras containing 5’ or 3’ half of E1 synonymous mutations from TC83 or 307537, or the central region of 306537 were constructed. **(B, C)** Viral replication of chimeric viruses infected at MOI 0.1. Supernatants were harvested at 12 **(B)** or 24 **(C)** hpi and infectious virus titered by focus forming assay (FFA). Each experiment was performed in triplicate three times independently and the mean and SD graphed. Statistical analysis was performed using an unpaired T-test. * >0.05, ** >0.001.

We hypothesized that altered macrophage replication fitness was driven by changes in RNA structure, which alter binding of RBPs to viral RNA. Therefore, to determine whether SNPs in the central region of E1 altered the underlying structure of E1, we performed in-cell SHAPE-MaP [34, 35] of cells infected with TC83 and TC83/E1_ID-syn_. Here, BHK cells were infected with TC83 or TC83/E1_ID-syn_ at an MOI of 0.1, treated with either DMSO (unreacted control) or the SHAPE chemical 1-methyl-7-nitroisatoic anhydride (1m7), and RNA lysates collected at 24hpi. SHAPE-MaP library preparation, sequencing, and analysis was performed as previously described [36], and SHAPE reactivity profiles generated for each viral genome (**Fig. 6A, Extended Data Fig. 6**). The SHAPE reactivity is indicative of the flexibility of each individual nucleotide, with low SHAPE reactivity correlating to paired nucleotides and high SHAPE reactivity correlating to unpaired nucleotides. Using these reactivity profiles as constraints for RNA folding, the secondary structure of TC83 and TC83/E1_ID-syn_ E1 was determined using RNAfold (**Fig. 6B, Extended Data Fig. 8).** Within the central region of E1, we observed conservation of several secondary structural elements (in grey) between TC83 and TC83/E1_ID-syn_. We also observed conservation of secondary structures in other regions of the viral genome, including the ribosomal frameshift motif in 6K/E1 (**Extended Data Fig. 5**). Notably, our data was found to be consistent with previously published SHAPE-MaP analysis of VEEV strain ZPC738 [37] (**Extended Data Fig. 5**) and we identified conserved secondary structures across all three viruses, lending further support to our findings. The central region of E1 responsible for the macrophage replication phenotype contains 11 SNPs (in blue). Three of these reside within the invariant RNA secondary structures identified (grey), and two SNPs (nts 10,481 and 10,633) were found to be unique to strain TC83 and another closely related IAB strain (AB66640; **Extended Data Fig. 7**). Of the remaining six SNPs, three were found within regions that displayed the most variable RNA secondary structure (nts 10,522, 10,606, 10,810; **Fig. 7A-C**, boxed base pairs). Since we hypothesized that changes in viral RNA structure contribute to emergence of epizootic VEEV in nature, we sought to determine whether SNPs were conserved across other epizootic or enzootic strains. We reasoned that RNA structures associated with epizootic emergence would not be unique to TC83 but would also be present in other epizootic isolates. To this end, we compared sequences across 29 epizootic strains (subtype IAB and IC) and 40 enzootic strains (subtype ID) (**Extended Data Fig. 7**). Indeed, with two exceptions, TC83 SNPs in the E1 central region were conserved across all IAB isolates. Phylogenetic analysis shows the presence of distinct lineages which largely correspond to distinct geographic distribution of these viruses[38–40]. Due to spatial evolution of these lineages, we speculated that epizootic-associated SNPs and RNA structures may also be lineage-specific, and not necessarily globally conserved across different lineages (compare epizootic sequences (red) within lineage K and Lineage L). Indeed, when we compared epizootic IAB (TC83; lineage L) and epizootic IC sequences (lineage K), we observed SNPs distinct to epizootic viruses versus enzootic within the one lineage, but which were different between epizootic viruses across distinct lineages. While almost all TC83-associated SNPs in this region were conserved across other IAB isolates, we observed that in IC isolates only SNPs at position 10,495 and adjacent to 10,522 (yellow highlight; **Extended Data Fig. 8**) differed between ID enzootic and IC epizootic isolates in this lineage (lineage K). This suggests that the evolutionary path to epizootic emergence is likely distinct to each outbreak and virus lineage. To determine whether epizootic sequences from different lineages adopt conserved RNA secondary structure despite the presence of distinct SNPs, consensus RNA structure predictions were generated for each epizootic and enzootic group from lineage M, L, and K using RNAalifold [17, 41] (**Extended Data Fig. 8D**). While overall the predicted secondary structure differed between all lineages, we observed that the 5’ structural element and first conserved structural element (**Fig. 6B**; highlighted in blue and grey in **Extended Data Fig. 8D**) were predicted to be conserved in both IAB and IC epizootic strains. This region encompasses four of the SNPs within the central E1 region responsible for differential macrophage replication. Collectively, this data shows that SNPs associated with macrophage replication fitness are conserved within lineages. Importantly, RNA structures are predicted to be conserved across epizootic viruses from distinct lineages despite variations in SNPs, suggesting that epizootic VEEV may evolve conserved RNA secondary structures that are functionally relevant for VEEV emergence.

**Figure 6.**
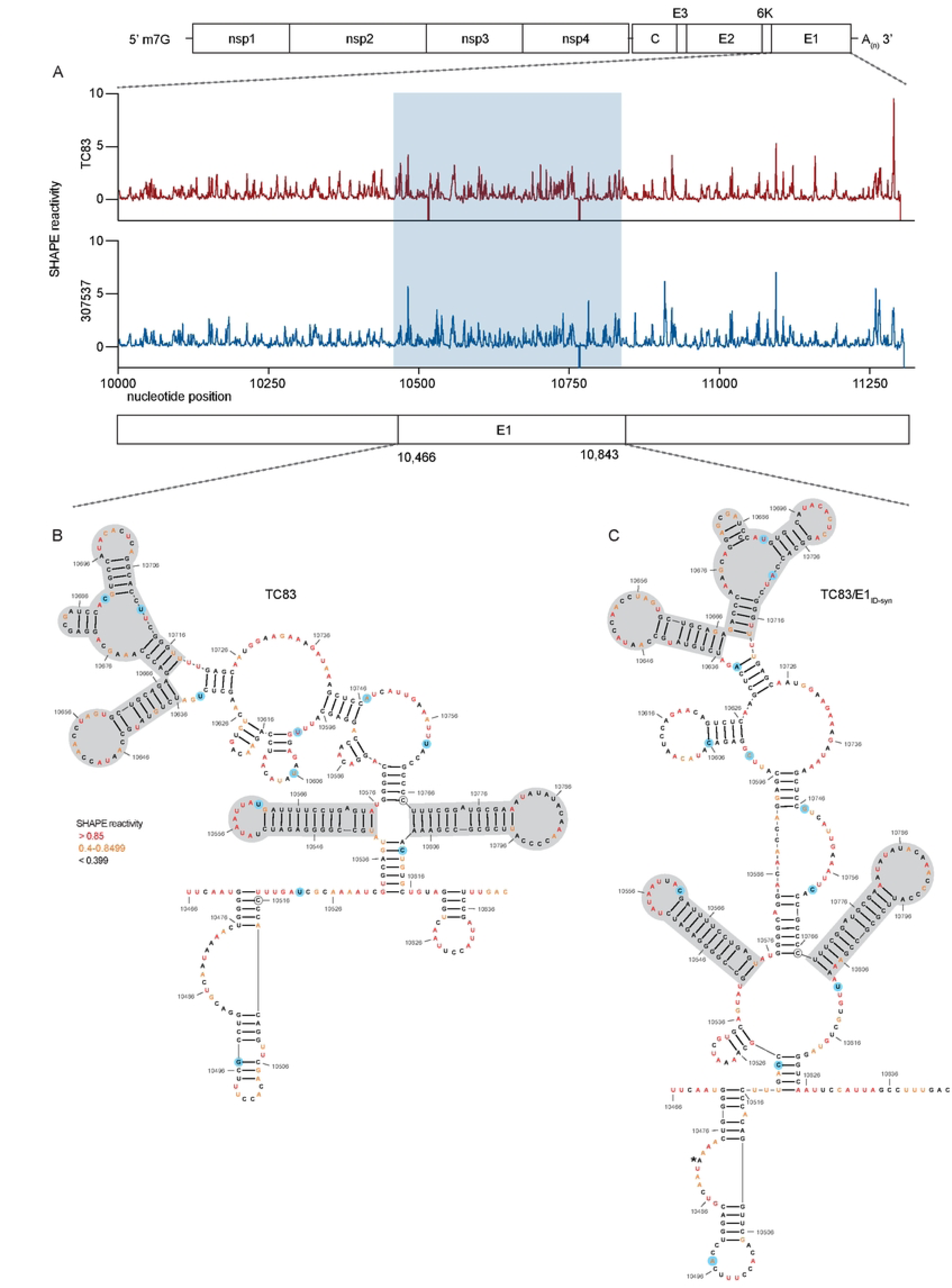
SHAPE-MaP analysis of TC83 and TC83/E1_IDsyn_ infected cells. RNA from BHK cells infected with TC83 or TC83/E1_IDsyn_ (MOI of 0.1) was analyzed by SHAPE-MaP. **(A)** Differential SHAPE reactivities of nucleotides in E1. The central region responsible for enhanced macrophage replication of TC83/E1_IDsyn_ is highlighted in blue. Secondary structures and SHAPE reactivities of nucleotides in the central domain of **(B)** TC83 and **(C)** TC83/E1_IDsyn_. Low reactive nucleotides (black) correspond to base-paired nucleotides and highly reactive nucleotides (orange, red) correspond to exposed bases. Structural elements conserved between both viruses are highlighted in grey, and SNPs are highlighted in blue.

## DISCUSSION

Repeated emergence of epidemic/epizootic VEEV as well as the emergence and re-emergence of other viral pathogens in recent times, has highlighted the need to better understand viral and host determinants that drive these processes. For VEEV, widespread vaccination of equines has been significant in the control of epidemic/epizootic outbreaks, though vaccines and therapeutics for use in humans remain a significant gap [42]. Moreover, the ability to produce significant viremia in humans, and the presence of susceptible urban mosquito vectors in VEEV endemic regions suggests significant potential for VEEV to evolve the ability to transmit in urban settings without the need for an equine amplification host. Thus, understanding how host and viral factors drive the evolution and emergence of VEEV and other viral pathogens in nature is paramount. While previous phylogenetic studies have emphasized the importance of amino acid mutations within E2 in the emergence of epizootic VEEV [9, 10, 43], our data supports an additional role for RNA structure in viral replication and cellular tropism, which has implications for immune evasion, dissemination, and transmission.

In this study we identified novel RNA structures that alter VEEV replication fitness specifically in macrophages, but not in other cell types. To our knowledge, this is the first time that VEEV RNA structure has been demonstrated to alter cellular tropism. In addition, we identified several RBPs which enhance replication of E1 enzootic mutants, namely FBL, DHX38, THRAP3, and UBAP2L. To our knowledge, none of these RBPs have previously been shown to play a role in facilitating alphavirus replication, or for the most part, other RNA viruses. Interestingly, FBL is well described to play a role in movement of plant viruses (reviewed in [44]) and has also been implicated in enhancing translation of structural genes of some viruses (reviewed in [45]). The mechanism of enhanced translation has been proposed to be mediated through protein-protein interactions, thus how FBL enhances enzootic VEEV replication through RNA interactions remains to be determined. As FBL is also known to play a role in processing and modification of rRNAs, recruitment to VEEV RNA may also impact cellular responses in a manner that benefits viral replication, analogous to what has previously been shown for other RBPs that are recruited to the SINV genome (e.g. HuR)[46]. Further RNA-protein interaction studies will reveal the molecular mechanisms by which these RBPs contribute to cell type specific replication fitness and tropism.

The observation that macrophage replication is specifically impacted by changes in E1 RNA is highly relevant, as myeloid cells are important targets early during in vivo infection and macrophages are important producers of IFN in this system [24, 25]. Notably, we observed that VEEV encoding E1 RNA sequences and RNA structures from an epizootic strain (TC83; IAB) replicated poorly in macrophages in relation to enzootic mutants. This is consistent with our prior studies, which similarly demonstrated that VEEV encoding either epizootic or enzootic 3’UTR sequences replicate differentially in an IFIT2-dependent manner [15]. While seemingly counterintuitive, we predict that diminished myeloid cell replication is associated with enhanced dissemination and viremia *in vivo*. In our proposed model (**Extended Data Fig. 9**), enhanced replication of enzootic mutants in myeloid cells in the lymph node leads to enhanced immune activation in neighboring cells which restricts viral replication in the periphery, leading to poor dissemination and viremia, reduced transmission, and possibly reduced pathogenesis. In contrast, epizootic mutants which replicate more poorly in these cells do not induce robust immune responses leading to more efficient dissemination and transmission. This model is also supported by studies with eastern equine encephalitis virus (EEEV), in which increased macrophage replication fitness leads to potent attenuation *in vivo* [24]. While increased macrophage replication with EEEV correlated with enhanced IFN production, we did not observe any significant difference in IFN expression or signaling between WT and E1 mutant VEEV. Nonetheless, we predict that multiple mechanisms (IFN-dependent and -independent) may possibly play a role in VEEV emergence, given the demonstrated role for this cytokine [12, 47].

In addition to the 3’UTR and E1, we are also examining how other VEEV RNA sequences and structures may contribute to myeloid cell replication fitness, and how this impacts dissemination, pathogenesis, and transmission. Based on our observations we propose a more complex mechanism of VEEV emergence which entails acquisition of multiple mutations across the genome that collectively facilitate viral entry, replication fitness, and immune evasion in amplification hosts and vector species that facilitate transmission during epizootic episodes. We predict that diminished macrophage replication fitness is a hallmark of epizootic VEEV isolates. Furthermore, we suggest that macrophage replication phenotypes may be a more accurate cell culture-based predictor of epizootic potential, instead of determination by E2 sequences alone. These findings highlight the complexity of factors that contribute to viral emergence and highlight the importance of examining multiple cell types and host factors.

## MATERIALS AND METHODS

### Cell lines

Vero C1008 and Raw264.7 cells were obtained from ATCC. All cell lines were maintained in DMEM supplemented with 10% heat-inactivated FBS (HyClone), 1% L-GlutaMAX (Gibco), and 1% nonessential amino acids (NEAA). Bone marrow derived macrophages (BMDMs) and Bone marrow derived dendritic cells (BMDCs) were generated independently from 10 to 20-week-old C57BL/6 mice. The mice were sacrificed, the femur and tibia were removed and cleaned. The bones were then briefly dipped in 70% EtOH to sterilized, followed by 1x PBS to remove any excess EtOH. The ends of the bones were then cut to expose the bone cavity and the bones were flushed with media using a 26.5G needle. The cells from one mouse were then divided over 3x 10cm non-tissue culture. To generate BMDMs, the dishes were grown in DMEM supplemented with 10% FBS (HyClone), 1% L-GlutaMAX (Gibco), 1% NEAA, 10,000 U/ml penicillin (Sigma), 10 mg/ml streptomycin (Sigma), and 20% L929-conditioned cell supernatant (described below). To generate BMDCs, the dishes were grown in DMEM supplemented 10% FBS (HyClone), 1% L-GlutaMAX (Gibco), 1% NEAA, 10,000 U/ml penicillin (Sigma), 10 mg/ml streptomycin (Sigma), 55mM β-mercaptoethanol and 20ng/ml GM-CSF (). On day 2 post harvesting, BMDMs were supplemented with 7ml BMDM media. On day 3, the cells were harvested by gently washing with PBS, followed by incubation with 10ml of 1mM EDTA in PBS for 5min at 37°C, and seeded for infection. On day 3 post harvesting, BMDCs were supplemented with 7ml BMDC media. On day 6, the non-adherent cells were harvested and seeded for infection.

The L929-conditioned cell supernatant was prepared by culturing L929 cells in a T175 until 90% confluence. This was then split into 6 new T175 flasks containing 45 ml of supplemented DMEM (10% GBS, 1% NEAA, 1% GluMAX) and cultured for 10 days at 37°C. Cell supernatants were then collected and centrifuged at 3000 rpm for 3 mins at 4°C. Lastly, supernatant was filtered using 45µM filter and stored at −20°C.

### Generation of Raw264.7 RIG-I^-/-^ and MDA5^-/-^ CRISPR cells

A doxycycline-inducible CRISPR/Cas9 expression vector (pSBtet-puro-Cas9-U6) was generated by cloning the Cas9-U6 portion of pX459 (Addgene #62988; [49] into pSBtet-pur (Addgene #60507; [50]). Cas9 was first cloned into pSBtet-pur using the following primers: Cas9.F: 5’-CATGAGACCGGTGCCACCATG-3’, Cas9.R: 5’-CATGAGGCGGCCGCCTACTTTTTCTTTTTTGCCTGGCCG, pSBtet-pur.F: 5’-CATGAG GCGGCCGCCTTCC-3’, pSBtet-pur.R: 5’-CATGAGACCGGTGGTGGCCGATATCTCAGAG. Post cloning, Cas9 was ligated into the pSBtet-pur backbone using the 5’ AgeI and 3’ Notl restriction sites. Following this, the U6 promoter was cloned into the new plasmid using the following primers: U6.F 5’-ACTACAGGTACC GAGGG-3’, U6.R 5’-TCAGTCCTAGGTCTAGAGC-3’, pSBtet-pur-Cas9.F 5’-TCAGTCCTAGGTCTAGAGC-3’, pSBtet-pur-Cas9.R 5’-ATGAAGGTACCACATTTGTAGAGGTTTTACTTGC-3’. U6 was ligated into pSBtet-pur-Cas9 using 5’ KpnI and 3’ AvrII restriction sites. As the new pSBtet-pur-Cas9-U6 plasmid contained an addition BbsI site, this was remove using site directed mutagenesis and the following primers: dBbsI.F 5’-TTGG GAAGAT AATAGCAG-3’, dBbsI.R 5’-CTGCTATTATCTTCCCAA-3’.

Sequence-specific gRNA sequences were designed using the Broad Institute Genetic Perturbation Platform gRNA design tool to target mouse Ddx58 and Ifih1. The primers detailed in **Table S1** were used to generate Dhx58 and Ifih1 gRNA oligonucleotides which were cloned into pSBtet-puro-Cas9-U6 as described previously [49].

Raw264.7 CRISPR cells were generated by electroporation of low passage Raw264.7 cells with Dhx58 and Ifih1 pSBtet-puro-Cas9-U6 using Amaxa Nucleofector II and Amaxa Cell Line Nucleofector Kit V (Lonza). Cells were selected with puromycin 3 days post-nucleofection, and Cas9/gRNA expression induced at 7 days post-nucleofection. Cells were treated for 14 days with doxycycline and KO efficiency of bulk cells validated using western blotting.

### Generation of full-length and recombinant viruses

Construction of the full length TC83 VEEV infectious clone has been described[16]. To introduce the E1 gene from KC344519 into TC83, a gBlock containing the E1 gene with flanking TC83 regions was generated (Table S2). The following primers were used to amplify two TC83 backbone fragments from the VEEV TC83 infectious clone described above: TC83 F1: 5’-GCTTGGTGCTGGCTACTATTG-3’, TC83 R1: 5’-CTCTTCGGATGCACCCTCAC −3’, TC83 F2: 5’-GATGCAGAGCTGGTGAG −3’, TC83 R2: 5’-GTTATACGAGATTCCCGCTTGG −3’. The backbone fragments were generated using Q5 high fidelity polymerase (NEB, M0491), treated overnight with DpnI and the DNA was purified using MicroElute Cycle-Pure Kit (Omega Bio-Tek). The fragments were assembled using Quantabio RepliQa HiFi assembly mix (#95190-D10) followed by transformation into NEB Stable Competent *E. coli*. Additional mutants were generated as follows. Fragments for mutant 1 were amplified using the following primers with corresponding plasmid: TC83/E1_ID-syn_ fw 5’-GCAAGATAGACAACGACG-3’ and rv 5’ GTCTCTGCAGCACTAGG 3’, TC83: fw 5’ CTGTATGCCAATACCAACC 3’ and rv 5’ CTGGCCCTTTCGTCTTC 3’. Mutant 2 fragments were generated using the same primers, but with the opposite plasmids. Fragments for mutant 3 were generated using the following primers: TC83/E1_ID-syn_ fw 5’ TTCAATGGGGTCAAAATAACTG 3’ and rv 5’ GTCAAAGGCTAATGGAATTGAC 3’, TC83 fw 5’GCAAGATAGACAACGACG 3’ and rv 5’ GGACCTGCAGTTATTTTGAC 3’, TC83 fw 5’ GTGCTGTAGGGTCAATTCC 3’ and rv 5’ CTGGCCCTTTCGTCTTC 3’. The fragments were generated and assembled as described above. Plasmids were linearized at MluI restriction sites located downstream of the poly(A) tail and genomic RNA was transcribed from the SP6 promoter in the presence of N7^m^G cap analog using the SP6 mMessage mMachine kit (Ambion). 1×10^7^ BHK21 cells were electroporated with approximately 2 µg of *in vitro* transcribed RNA using a GenePulser Xcell electroporator (Bio-Rad) to generate P0 virus stocks.

### Focus-forming assays

Vero E6 monolayers were infected with serial 10-fold dilutions of infectious samples for 1 hour at 37°C, then overlaid with 100 µl per well of medium (0.5x DMEM, 5% FBS) containing 1% carboxymethylcellulose, and incubated for 20 to 22 hours at 37°C with 5% CO_2_. Cells were then fixed by adding 100 µl per well of 2% paraformaldehyde directly onto the overlay at RT for 2 hours. After removal of overlay media and fixative, cells were washed 3x with PBS and incubated with antibodies specific for VEEV E2 glycoprotein (gift of Dr. Michael Diamond) for 2 hours at RT in FFA permeabilization buffer (1x PBS, 0.1% saponin, and 0.1% BSA). Mouse anti-VEEV E2 (clone 36.E5) were produced and purified from a clonal hybridoma cell line and which was a generous gift from Dr. Michael Diamond (Washington University School of Medicine, St Louis). Cells were washed 3x in ELISA wash buffer (1x PBS, 0.05% triton X-100), then incubated with species-specific HRP-conjugated secondary antibodies (Sigma and ThermoFisher) for 1 hour at RT in FFA permeabilization buffer. Monolayers were washed 3x with ELISA buffer and foci were developed by incubating in 50 µl/well of TrueBlue peroxidase substrate (KPL) for 5 to 10 minutes at RT, after which time cells were washed twice in water. Well images were captured using Immuno Capture software (Cell Technology Ltd.), and foci counted using BioSpot software (Cell Technology Ltd.). All samples were titered in duplicate and calculated titers averaged for each duplicate.

### Viral growth kinetic assays

Multistep viral growth kinetics were performed by infecting Raw264.7 with WT or mutant VEEV TC83 viruses at a MOI of 0.1. Cells were seeded 18-20hrs prior to infection. Viral titers were determined for indicated time points post-infection by removing cell culture supernatant, replacing it with fresh growth media, and subsequently measuring viral titers through FFA. All experiments were performed three or four times independently in triplicate. Statistical analysis was performed by calculating area under the curve (AUC) and performing unpaired t-test on AUC values calculated for each experiment. P values are reported in each figure.

### siRNA knock-down

DsiRNA transfections were done in 96 well format. DsiRNAs used are listed in Table S3. Transfection mix was made up of 10nM DsiRNA pool, 0.2µl *Trans*IT-X2 Dynamic Delivery System (Mirus, 6003) and supplement-free DMEM and incubated at RT for 20mins. 2E4 Raw264.7 cells were combined with the transfection complexes and seeded into a 96 well plate. 24 hours post transfection were mock infected or infected with either TC83 or TC83/E1_ID-syn_ at an MOI of 0.1. 24 hours post infection, supernatant was collected and titered as describe previously. Cell viability was then assess using alamarBlue Cell Viability Reagent (Invitrogen, DAL1025) as described by the manufacturer.

### IFNAR blocking antibody infections

Raw264.7 cells were seeded 24h prior to infection. One hour prior to infection, the cells were pretreated with 10µg of mouse IgG2a isotype control (InVivoMAb, BE0085) or IFNAR1 Monoclonal Antibody (MAR1-5A3, Invitrogen 16-5945-85), infected with TC83 or TC83/E1_ID-syn_ at an MOI of 0.1 in the presence of antibody. Infectious virus from cell culture supernatants harvested at 10 and 22 hpi was titered by FFA. Each experiment was performed three times independently. Cell lysates were collected at 22hpi for RT-qPCR analysis.

### RT-qPCR

Cell lysates were prepared using Quick-RNA MiniPrep Kit (Zymo Research, Cat# 11-328) according to manufacturer’s protocol. Samples were DNase I (NEB, M0303) treated for 20 mins at 37°C, followed by inactivation of DNase I in 0.1M EDTA for 10 mins at 70 °C. cDNA was generated with 100ng/10µl reaction using iScript cDNA synthesis kit (Bio-rad, 1708890). qPCR was then run with 1µl of cDNA using iTaq Universal Probes Supermix (Bio-rad, 1725130) on Bio-Rad CFX96 Real-Time System. The following primer probe assays were used: Ifit1 (IDT, Mm.PT.58.32674307), IFN-beta (IDT, Mm.PT.58.30132453.g), ISG15 (IDT, Mm.PT.58.41476392.g) and VEEVset3 (nt9835-9856) (IDT, probe sequence: /56-FAM/TTT GTC TGG /ZEN/CTG TGC TTT GCT GC/3IABkFQ/).

### Western blotting

Cell lysates were generated by washing monolayers with PBS followed by incubation with RIPA lysis buffer (Thermofisher, cat# 89901) supplemented with Halt protease inhibitor cocktail (Thermofisher, cat# 78429) on ice for 5 min. Lysates were then scraped, transferred to microcentrifuge tubes, pulse vortexed and further incubated on ice for 15 mins. Hereafter, lysates were centrifuged at 16,000xg for 20 mins at 4°C and supernatants were transferred to a new tube. Proteins were separated by on a 4-20% Mini-PROTEAN TGX precast protein gel (Bio-rad), transferred to a nitrocellulose membrane (Amersham, 10600008), and then labeled for proteins. The following antibodies were used: beta-Actin Mouse mAb (Cell signaling, 8H10D10), beta-Actin Rabbit mAb (Cell signaling, 13E5), Rig-I mAb (Cell signaling, D1466), MDA-5 Rabbit mAb (Cell signaling, D74E4), Fibrillarin/U3 RNP Rabbit pAb (ABclonal, A1136), Dhx38 (ABclonal A4341), Goat-anti-Rabbit IRDye 800 (Licor, 926-32211), Goat-anti-mouse IRDye 680 (Licor, 926-68070).

### Immunoprecipitation-mass spectrometry

Cell lysates were generated from Raw264.7 cells by resuspending cells in 1X CHAPS lysis buffer (10mM HEPES, 200mM NaCl, 1% CHAPS, 10mM MgCl2, protease inhibitor (Thermo Scientific Pierce, PIA32955), 200U/ml murine RNase inhibitor (NEB, M0314)). Lysates were then passed through a 25G needle 4x and incubated on ice for 15min to ensure lysis. Thereafter, lysates were centrifuged at 16,000xg for 20 mins at 4°C and supernatants were transferred to a new tube. 50µl of Dynabeads protein G (Thermo, #10003D) were washed 2x in 500µl lysis buffer, after which they were incubated in 200µl lysis buffer along with 12µg of mouse J2 IgG2a Or mouse IgG2a isotype control (InVivoMAb, BE0085) for 30 mins at RT. Beads were then washed 3x in lysis buffer, incubated with 3mg of Raw264.7 lysate on the rotator for 2hrs at RT, washed again 3x in lysis buffer followed by 3x with freshly prepared 20mM ammonium bicarbonate. Samples were then trypsin digested in 20µl 20mM ammonium bicarbonate and incubated with 10µl of 10ng/µl sequencing grade trypsin (Promega, cat# V5111) at 37°C for 3hrs at 1500rpm. The supernatant was then carefully removed from the beads and the beads where washed 2x in 30µl 20mM ammonium bicarbonate, and all fractions were pooled. Samples were then reduced by adding tris(2-carboxyethyl)phosphine (TCEP) to a final concentration of 1mM and incubated at 37°C for 1h. Freshly prepared iodoacetamide (Thermo, cat# 90034) was then added to a final concentration of 10mM and incubated at RT for 30min in the dark, followed by quenching with final concentration 2mM N-AcetylCysteine. Samples were then cleaned-up and concentrated using C18 columns (Thermo-Pierce, cat# 89870) according to the manufacturers protocol. After clean-up, formic acid was added to the samples at a final concentration of 0.1%. Samples were analyzed by LC/MS at University of Washington’s Proteomics Resource (UWPR).

### SHAPE-MaP

VERO cells were seeded at 25E6 cells per 10cm dish. 24 hours post seeding, cells were infected with either TC83 or TC83/E1_ID-syn_ at MOI 0.1. At 24hpi culture media was aspirated and cells were washed once with 1x PBS. In-cell SHAPE modifications were made by adding fresh 500µl of 100mM 1-Methyl-7-nitroisatoic anhydride (1M7) (Sigma-Aldrich, 908401) in DMSO to 4.5ml pre-warmed culture media to the dish and incubated for 3min at 37C. This was repeated 3x to increase modifications. Unmodified samples were similarly treated with DMSO. After treatment, whole-cell RNA was purified using TRIzol reagent (Fisher Scientific) according to the manufacturers protocol. Samples were treated with TURBO DNase (Thermo Fishter, AM2238) for 30min at 37C to remove any DNA. Polyadenylated RNA was then isolated from the whole-cell RNA using NEB Oligo d(T)25 Magnetic beads (NEB, S1419S) according to the manufacturers protocol. To generate the denatured controls, 1µg of TC83 DMSO and TC83/E1_ID-syn_ DMSO polyA purified RNA were heated to 95C for 2min in 1x DC buffer (50mM HEPES (pH 8.0), 4mM EDTA) with an equal volume of 100% formamide. Samples were then immediately transferred to a new tube containing fresh 1M7 to a final concentration of 10mM and heated at 95C for 2min, after which the samples were placed on ice. DC control RNA was then purified using G-50 columns (GE healthcare, 25-5330-01). All samples were then prepared for sequencing using the randomer library prep workflow protocol described in Smola et al. [36]. Samples were sequenced by Illumina NGS at the Fred Hutch Cancer Center genomics core. Sequencing data was analyzed using Shapemapper2 as previously described.

## ACKNOWLEDGEMENTS

NIH grant R01 AI155416 and UW Royalty Research Fund A172539 supported this work. This work is supported in part by the University of Washington’s Proteomics Resource (UWPR95794) and the Genomics & Bioinformatics Shared Resource, RRID:SCR_022606, of the Fred Hutch/University of Washington/Seattle Children’s Cancer Consortium (P30 CA015704). The authors thank M.S. Diamond and I. Frolov for their generosity with VEEV reagents. The authors have no financial conflicts to disclose. The data reported in this manuscript are tabulated in the main paper and in the supplementary materials.

---

**Extended Data Figure 1.**
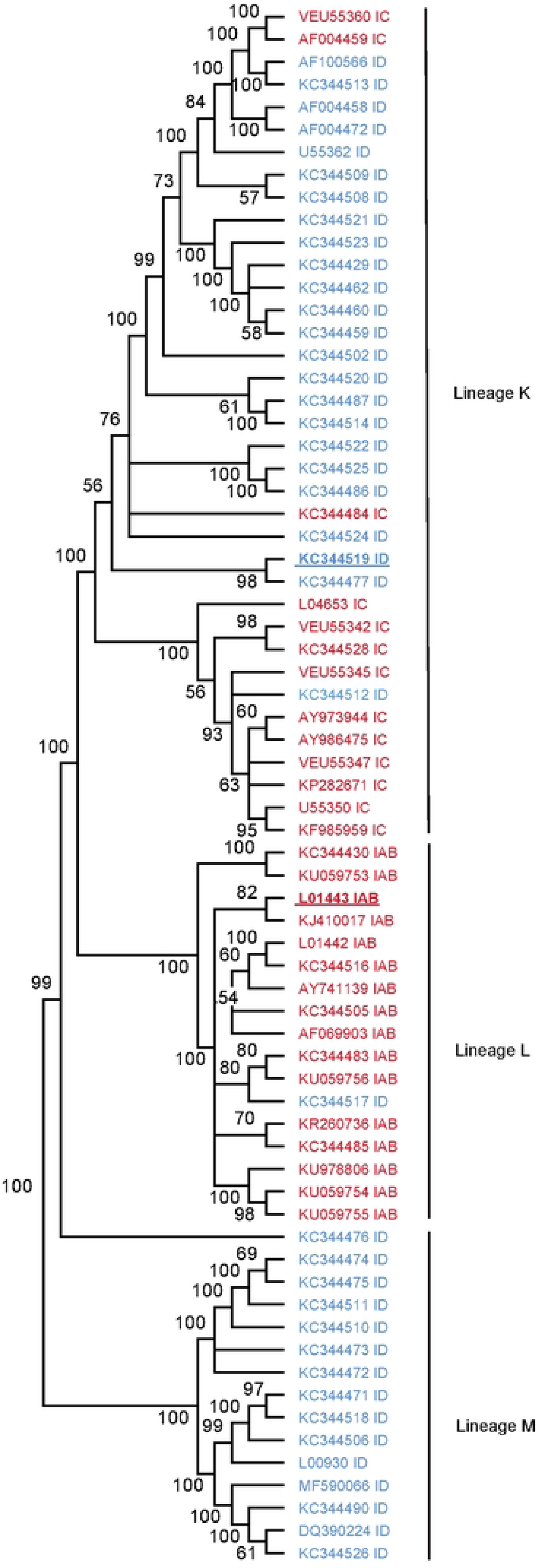
Phylogenetic tree of VEEV IAB, IC and ID subtypes. The optimal phylogenetic tree of lineages K, L and M (previously described in [39]) as determined by the neighborhood-joining method [48]. Shown next to each branch is the percentage of replicate trees in which the associated taxa clustered together in the bootstrap test (1000 replicates).

**Extended Data Figure 2.**
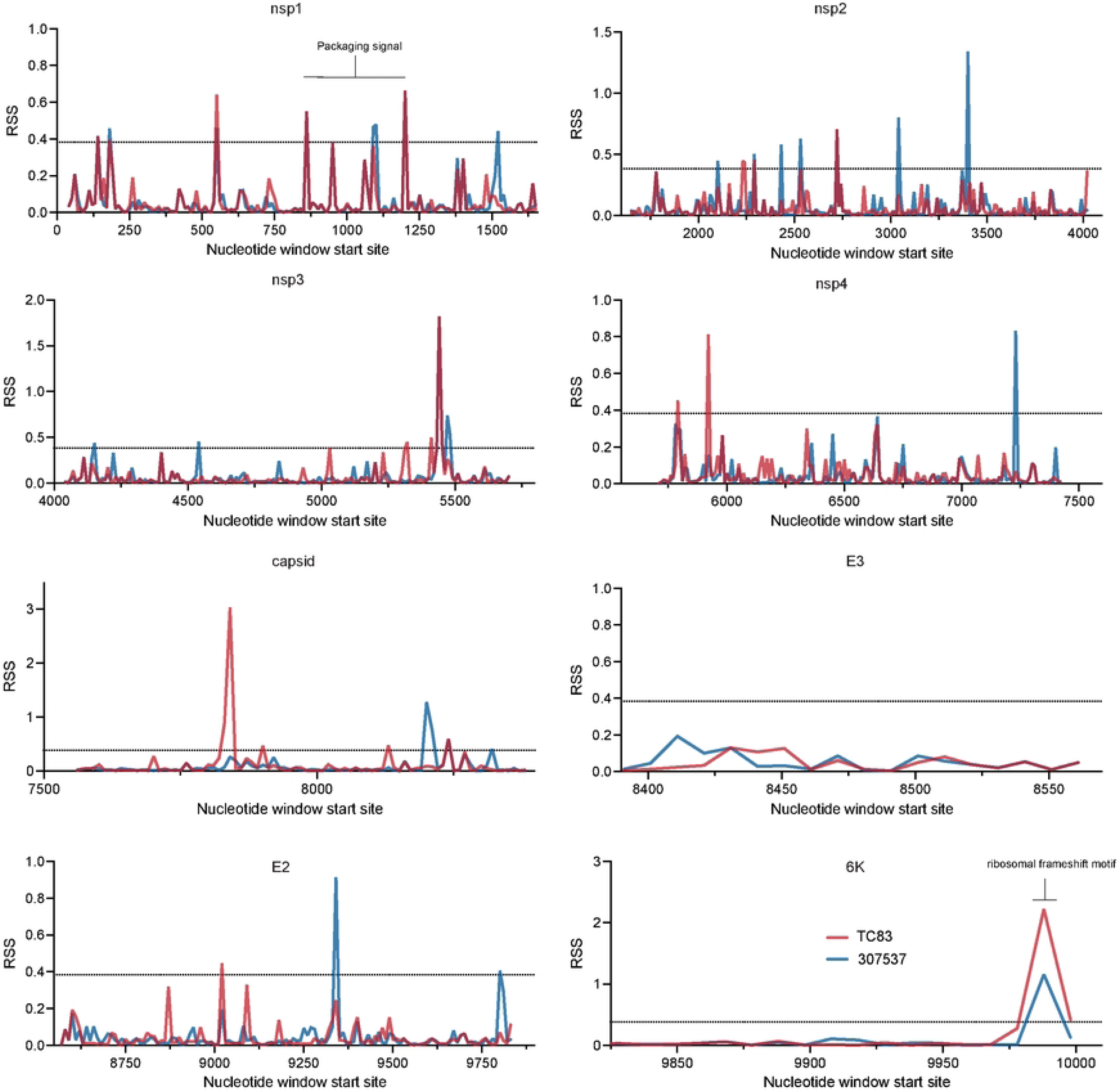
Sliding window analysis of the relative structure score (RSS) across the VEEV genome broken up by gene. The RSS was calculated as the minimum free energy (MFE)/ensemble diversity for each window of 50 nucleotides with a step size of 10. TC83 is shown in red and 307537 is shown in blue. Two standard deviations from the mean was calculated across the entire genome and is represented as a dotted line.

**Extended Data Figure 3.**
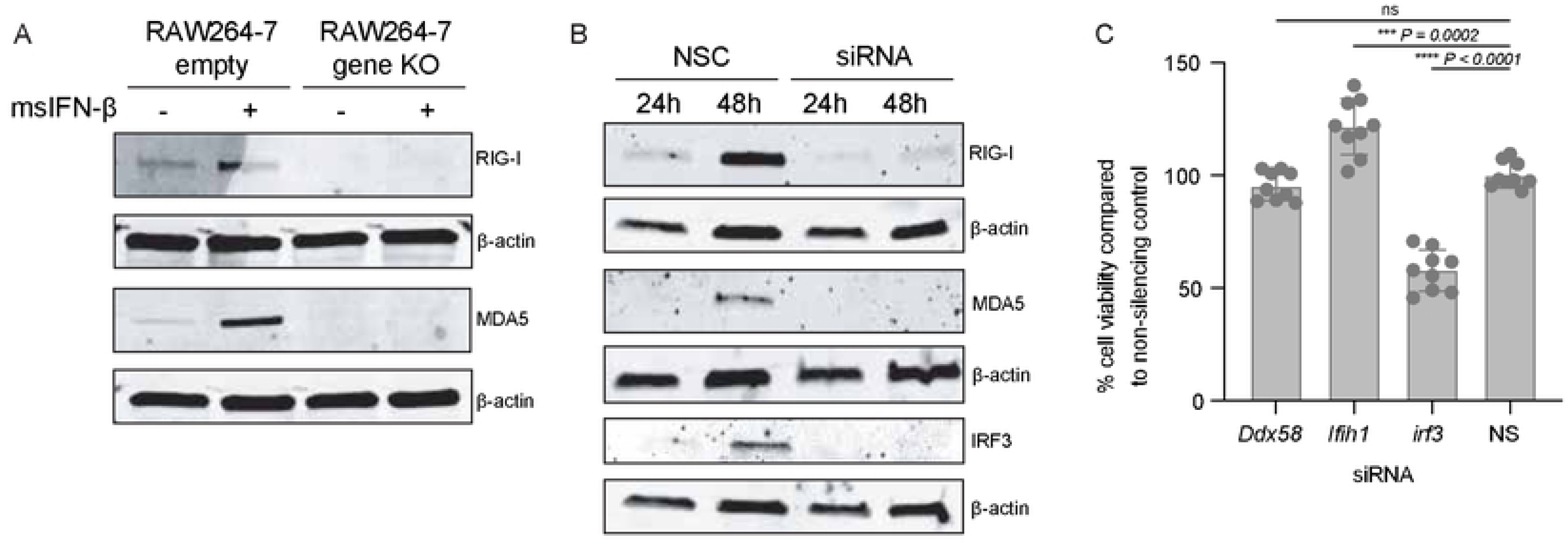
Validation of CRISPR KO and siRNA KD in Raw 264.7 macrophages. **(A)** Western blots from Raw264.7 6hrs after treatment +/− 100U/ml msIFN-β. **(B)** Western blot of Raw264-7 after transfection with NSC or protein of interest siRNA siRNA pool (10µM) pool for 24hrs or 48hrs. **(C)** Cell viability was determined using alamarblue Cell Viability Reagent and calculated as a percentage compared to the NSC. Statistical analysis was performed using GraphPad Prism 9, using an unpaired T-test.

**Extended Data Figure 4.**
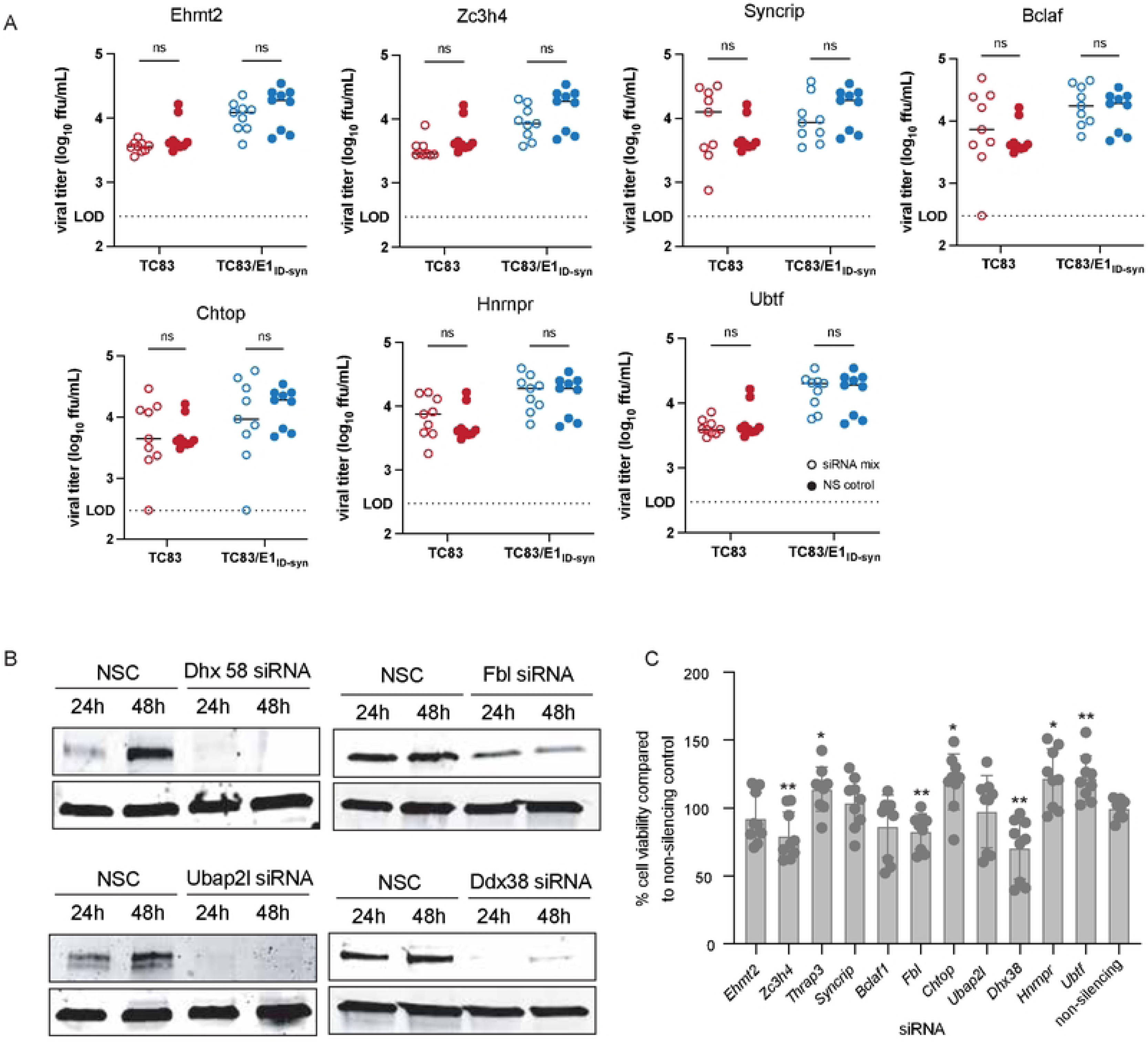
Validation of additional mass spectrometry hits. To evaluate the effect of these genes on viral replication **(A)** Raw264-7 cells were transfected with 10mM of a pool of 3 siRNA targeting proteins of interest for 24hrs, after which they were infected with TC83 or TC83/E1_ID-syn_. Supernatants were collected at 24hpi and infectious virus was titered using FFA. For visual clarity, the individual siRNAs along with the non-silencing control (NSC) are graphed individually, however the NSC is the same in all graphs. Each experiment was performed in triplicate three times independently and the mean and SD are graphed. **(B)** Western blot of Raw264-7 transfected for 24h or 48h with NSC or protein of interest siRNA pool (10µM). **(C)** Cell viability was determined using alamarblue Cell Viability Reagent and calculated as a percentage compared to the NSC. Statistical analysis was performed using GraphPad Prism 9, using an unpaired T-test. * >0.05, **>0.001.

**Extended Data Figure 5.**
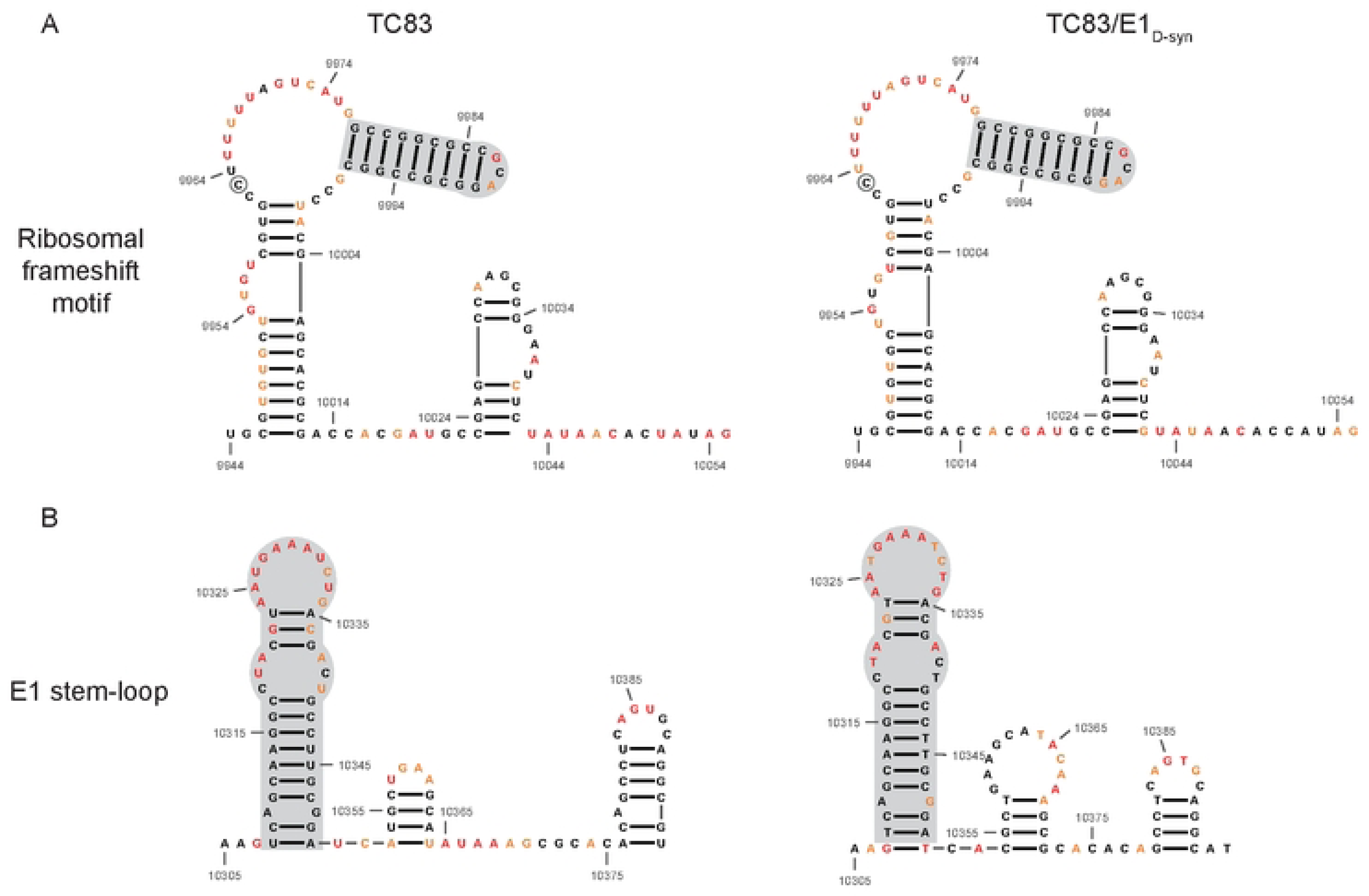
Previously described stable RNA structures conserved in SHAPE-MaP informed RNA secondary structure. Previous work by Kutchko *et al.* [37] performed *in vitro* SHAPE-MaP of the enzootic ID VEEV strain, ZPC738, and identified stable RNA structures across the VEEV genome. Displayed here are the SHAPE-MaP informed secondary structure predictions of **(A)** the ribosomal frameshift motif and **(B)** an E1 stem-loop for TC83 and TC83/E1_ID-syn_. Shaded in grey are the conserved regions identified between the previously described stable structures in ZPC738 and the *in vivo* SHAPE-MaP data generated for TC83 and TC83/E1_ID-syn_.

**Extended Data Figure 6.**
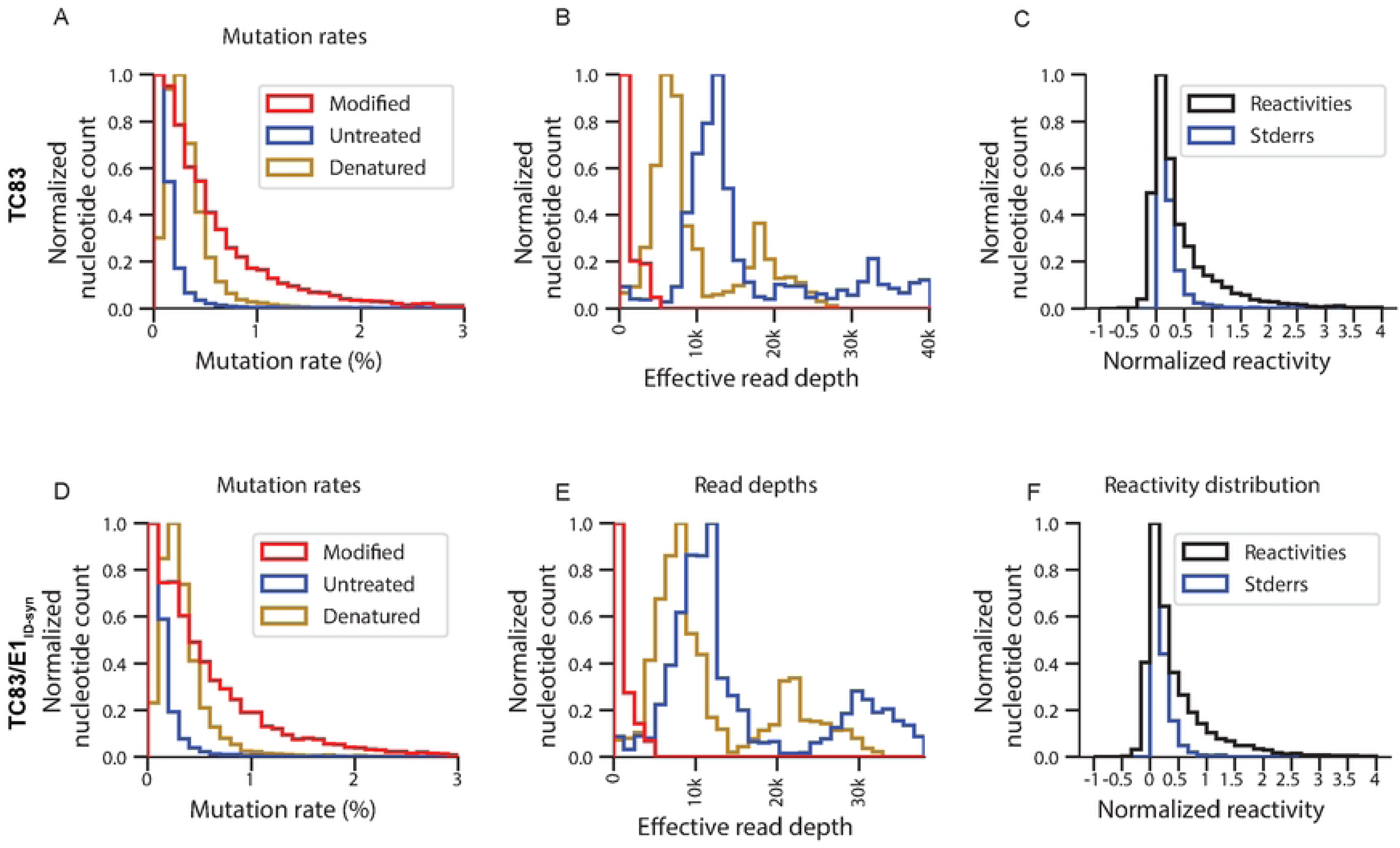
Quality matrixes for SHAPE-MaP for TC83 and TC83/E1_IDsyn_. Mutation rates for modified, untreated and denatured control **(A)** TC83 and D. TC83/E1_ID-syn_. Read depths for modified, untreated and denatured control **(B)** TC83 and E. TC83/E1_ID-syn_. The distribution of the SHAPE-MaP reactivities and the standard error of the reads for C. TC83 and F. TC83/E1_ID-syn_.

**Extended Data Figure 7.**
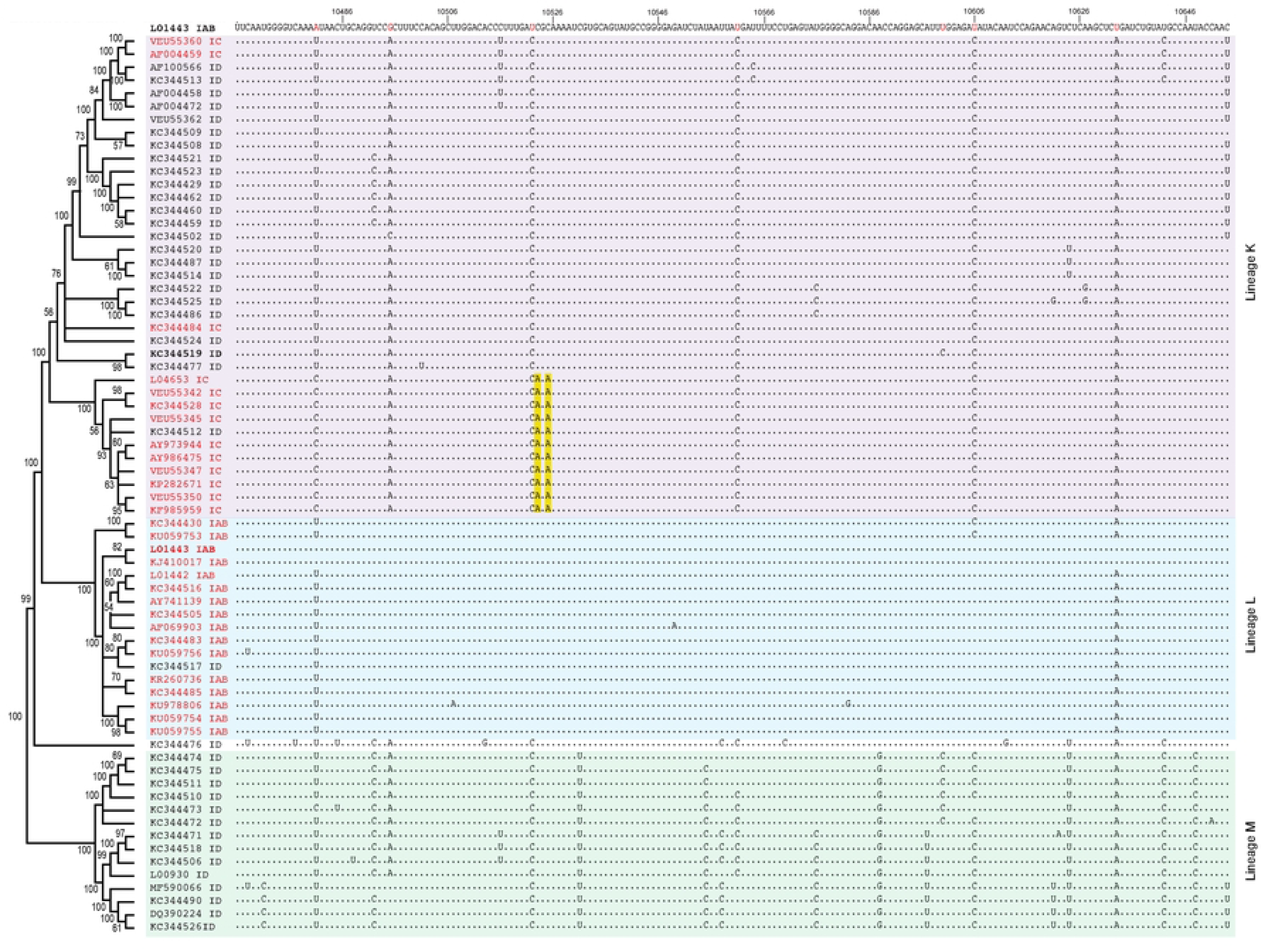
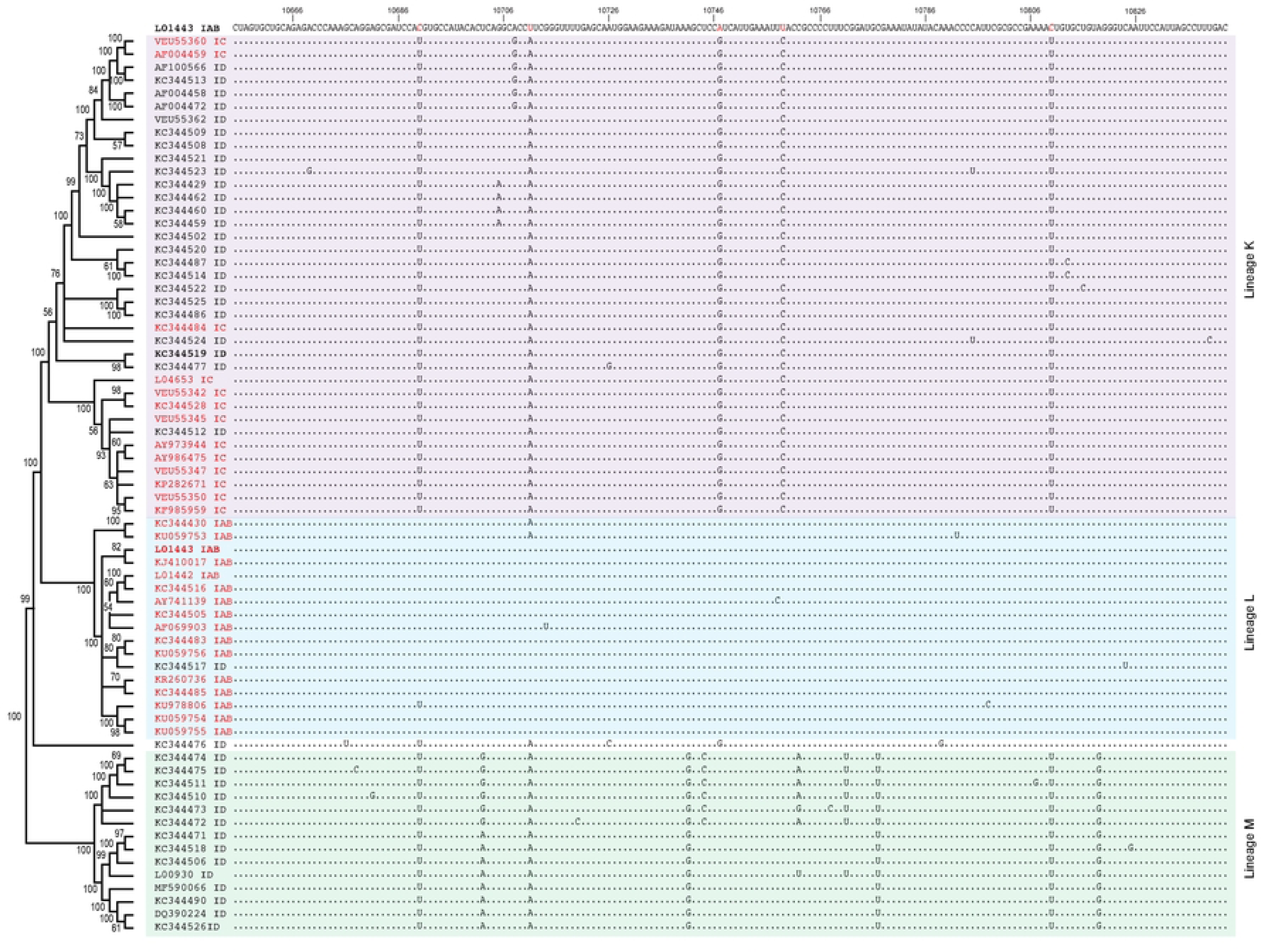
Sequence alignment of the core E1 region. Alignment contains VEEV sequences from lineage K, L and M shown in the phylogenetic order determined in supp figure 1. Lineage K sequences are shaded in purple, lineage L sequences are shaded in blue and lineage M sequences are shaded in green. The alignment was made using TC83 (L01443 IAB) as the reference sequence. Varying nucleotides between TC83 and 307537 (KC344519 ID) are highlighted in red in the reference sequence. Identical nucleotides are represented as periods (.).

**Extended Data Figure 8.**
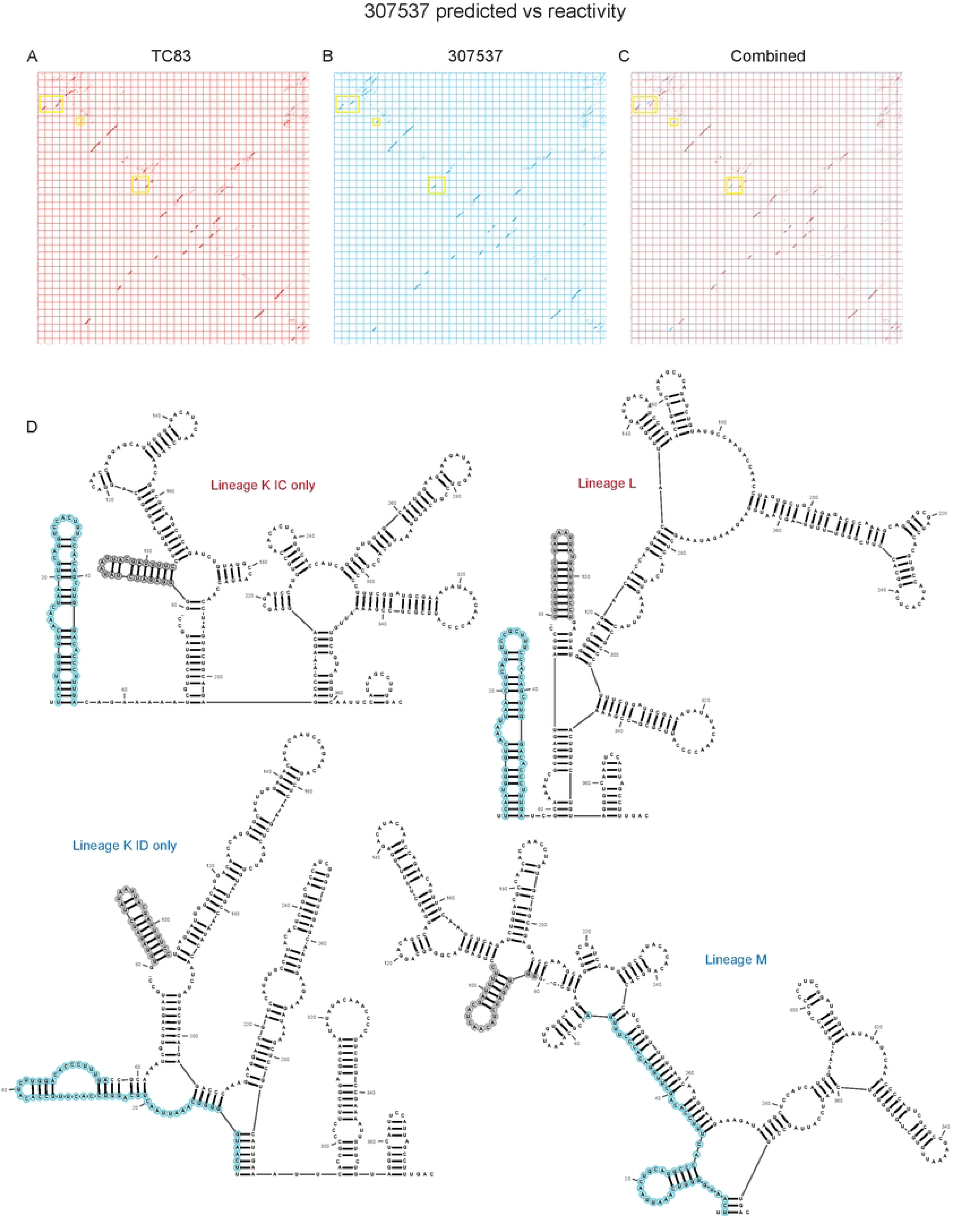
TC83 E1 SNPs are conserved in other epizootic strains and are lineage specific. Dot plots from RNAfold [17] predictions of individual SNPs within the E1 core region (10,466-10,843) for **(A)** TC83, **(B)** TC83/E1_ID-syn_ and **(C)** overlayed dotblots. Yellow boxes highlight regions with differences in RNA structure predictions. **(D)** Predicted RNA secondary structures from RNA alignfold [41]of sequences from lineages K (divided into epizootic IC and enzootic ID), L and M. Steml-oop conserved in epizootic lineages is highlighted in blue, and stem-loop conserved in all lineage K and L highlighted in grey.

**Extended Data Figure 10.**
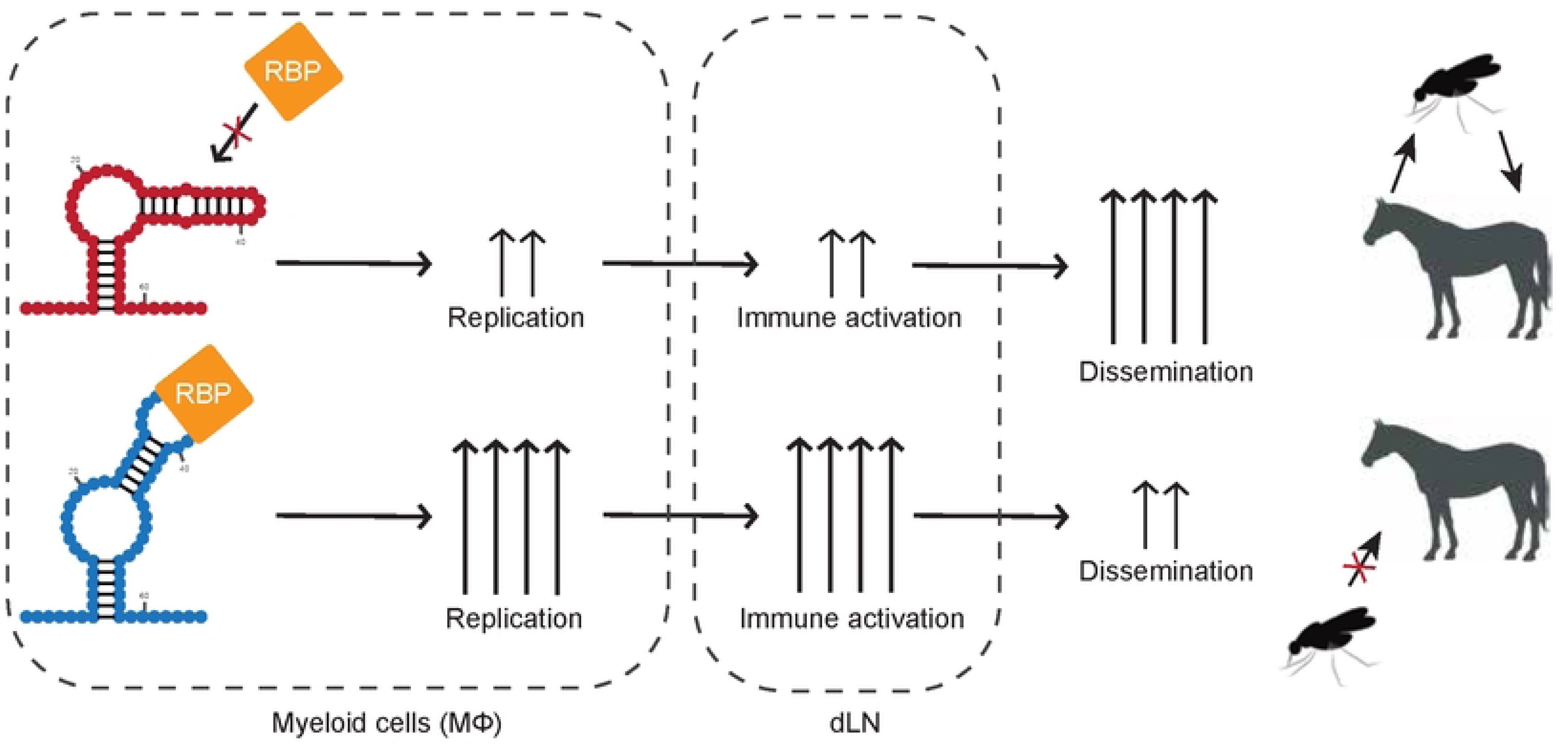
Model of RNA structure contributions to emergence and pathogenesis of VEEV. Following infection and trafficking to the proximal draining lymph node of an infected host, enzootic E1 RNA structures lead to recruitment of proviral factors (IFN-independent RBPs) which enhance replication specifically in macrophages, and possibly other myeloid cell types. Enhanced viral replication in the lymph node leads to increased accumulation of dsRNA and stimulates antiviral responses in host cells, preventing efficient dissemination and viremia. Reduced viremia leads to reduced transmission and possibly reduced pathogenesis.

**Table S1:**
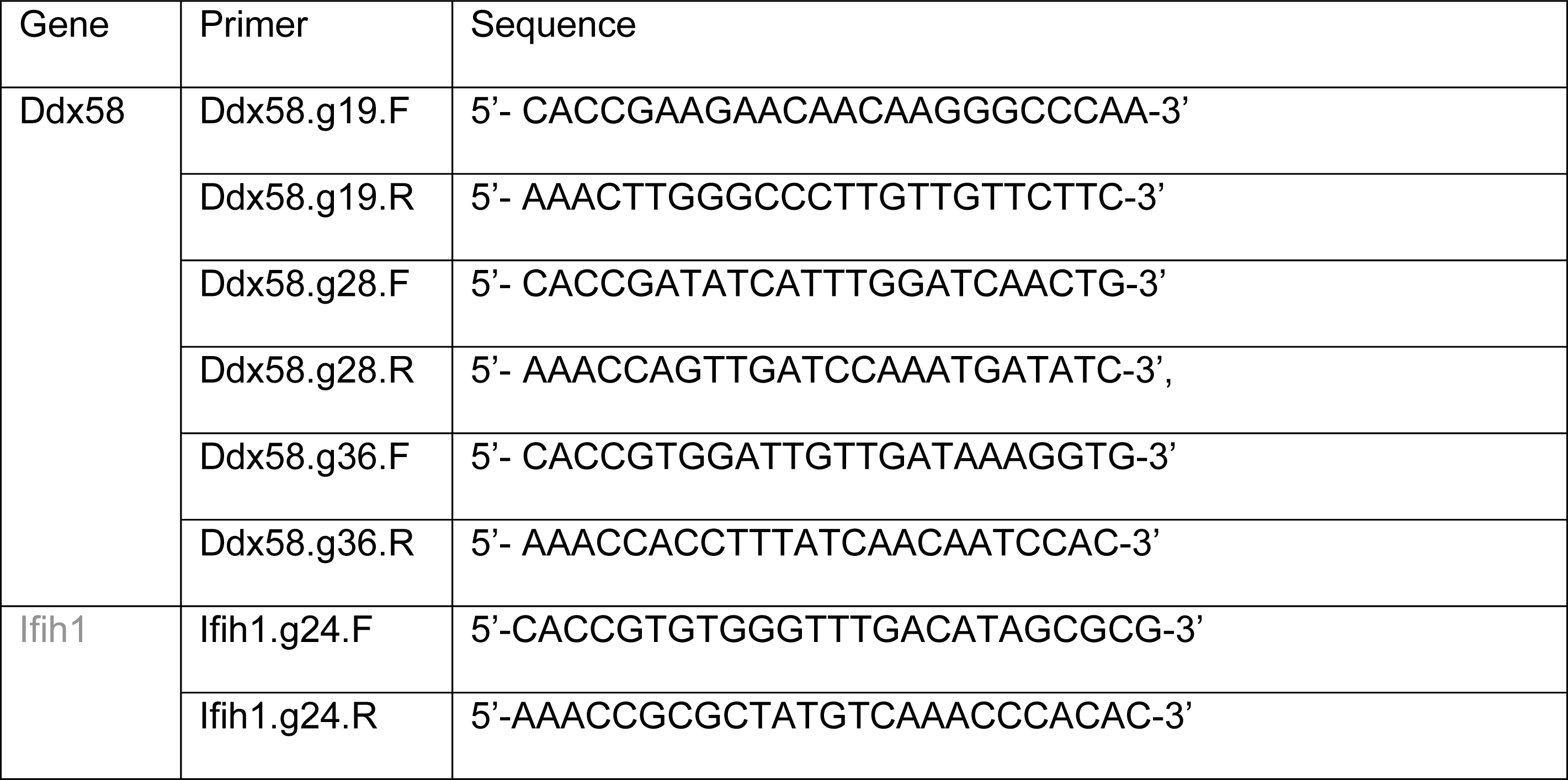
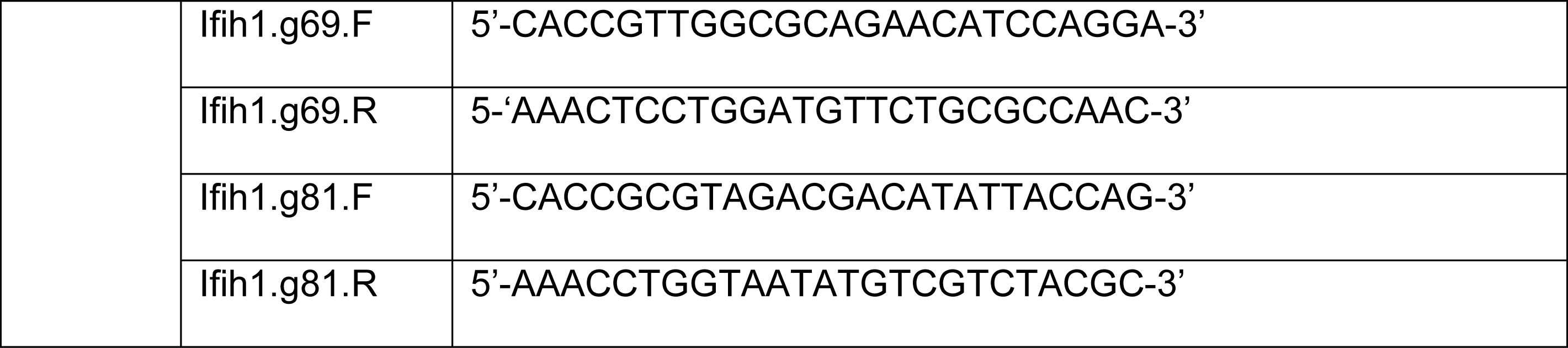
Primer sequences for generation of RIG-I and MDA5 CRISPR cell lines.

**Table S2:**
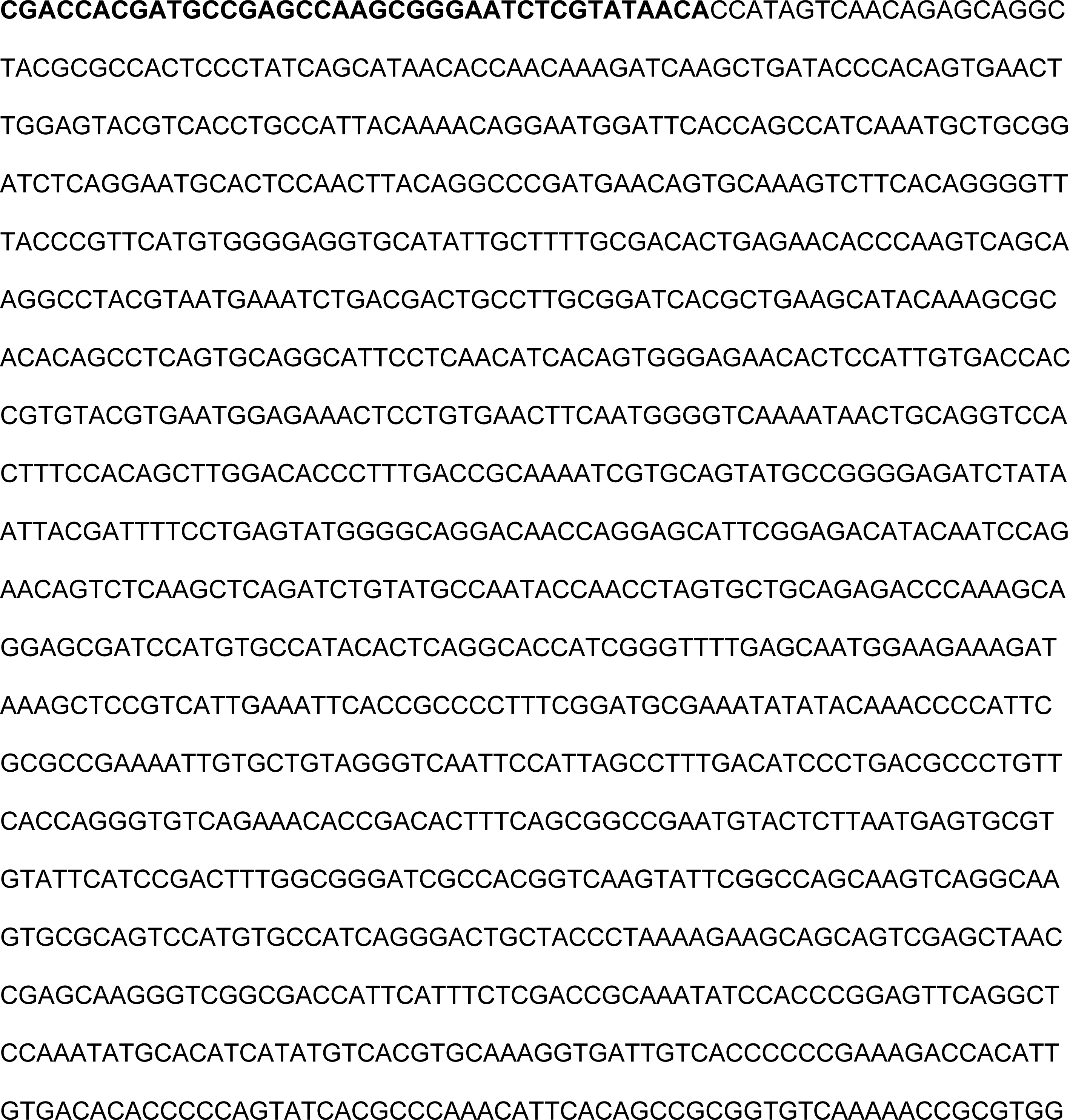
KC344519 E1 gene block.

**Table S3.**
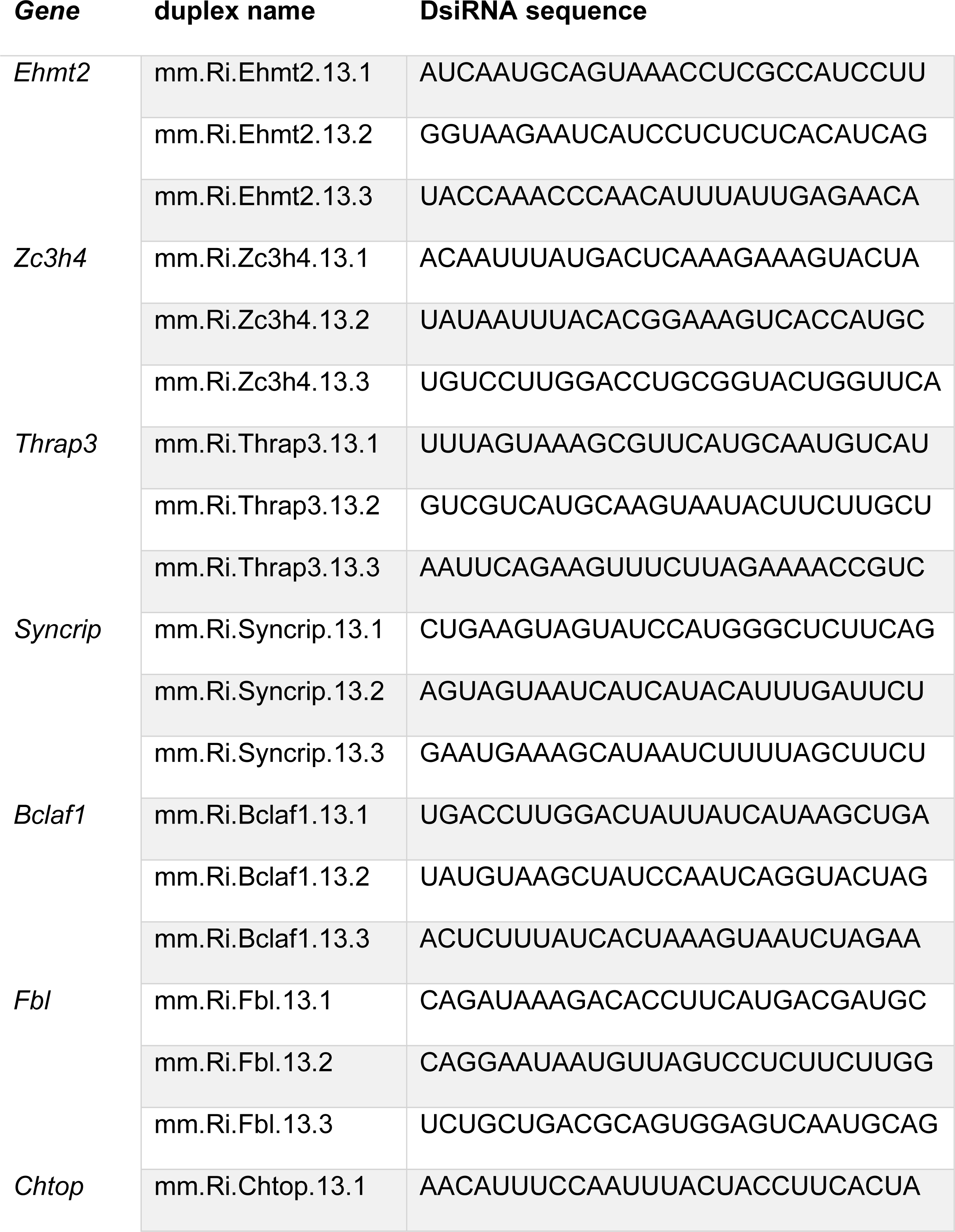
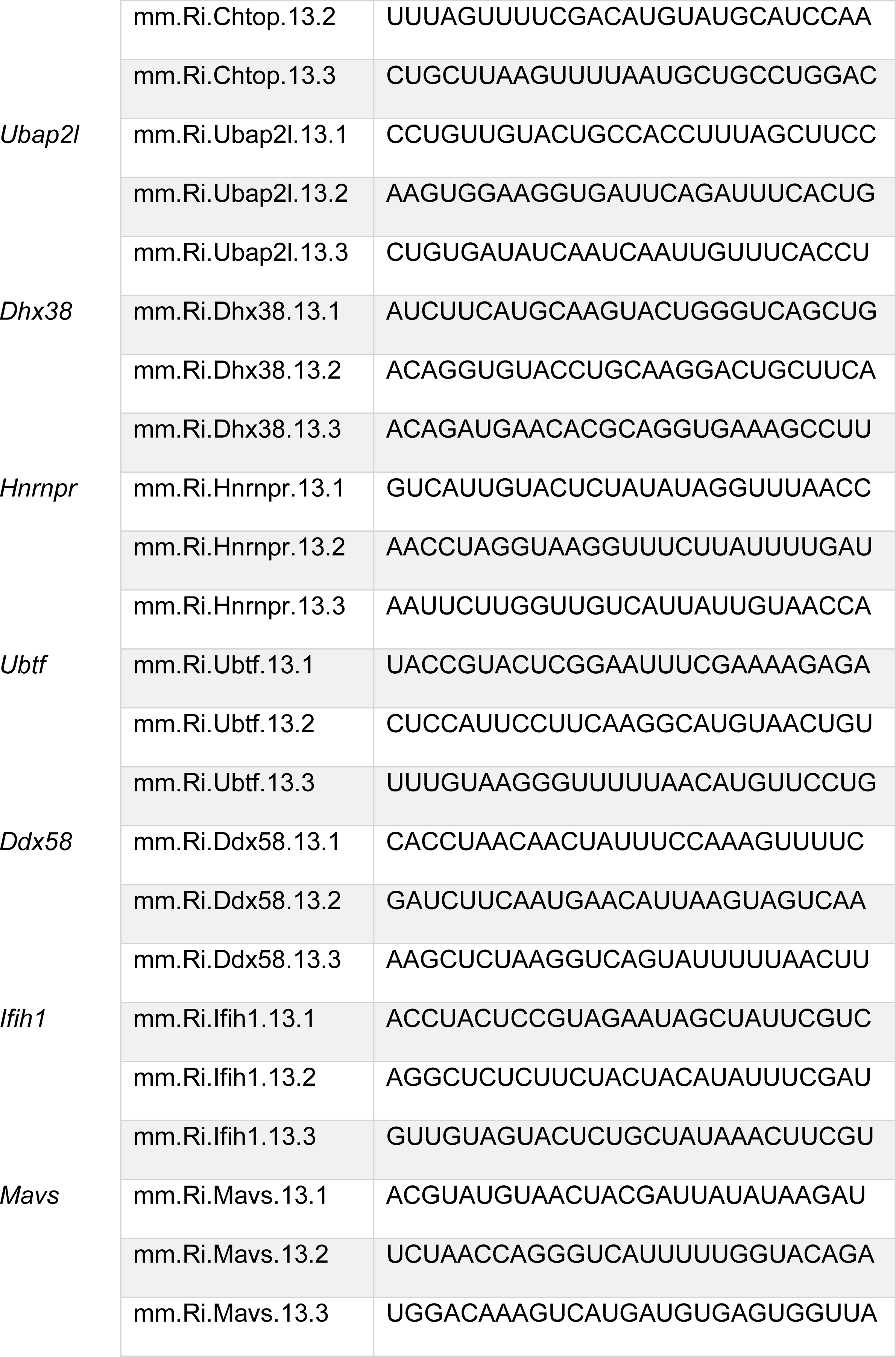
DsiRNA sequences.

